# CK2 alpha prime and alpha-synuclein pathogenic functional interaction mediates synaptic dysregulation in Huntington’s disease

**DOI:** 10.1101/2020.10.29.359380

**Authors:** Dahyun Yu, Nicole Zarate, Angel White, De’jah Coates, Wei Tsai, Carmen Nanclares, Francesco Cuccu, Johnny S. Yue, Taylor G. Brown, Rachel Mansky, Kevin Jiang, Hyuck Kim, Tessa Nichols-Meade, Sarah N. Larson, Katie Gundry, Ying Zhang, Cristina Tomas-Zapico, Jose J. Lucas, Michael Benneyworth, Gülin Öz, Marija Cvetanovic, Alfonso Araque, Rocio Gomez-Pastor

## Abstract

**Background:** Huntington’s Disease (HD) is a neurodegenerative disorder caused by a CAG trinucleotide repeat expansion in *the HTT* gene for which no therapies are available. This mutation causes HTT protein misfolding and aggregation, preferentially affecting medium spiny neurons (MSNs) of the basal ganglia. Transcriptional perturbations in synaptic genes and neuroinflammation are key processes that precede MSN dysfunction and motor symptom onset. Understanding the interplay between these processes is crucial to develop effective therapeutic strategies to treat HD. We investigated whether protein kinase CK2α’, a kinase upregulated in MSNs in HD and previously associated with Parkinson’s disease (PD), participates in the regulation of neuroinflammation and synaptic function during HD progression.

**Methods:** We used the heterozygous knock-in zQ175 HD mouse model and compared that to zQ175 mice lacking one allele of CK2α’. We performed neuropathological analyses using immunohistochemistry, cytokine proteome profiling, RNA-seq analyses in the striatum, electrophysiological recordings, and behavioral analyses. We also used the murine immortalized striatal cell lines ST*Hdh*^Q7^ and ST*Hdh*^Q111^ and studied the expression of various synaptic genes dysregulated by CK2α’.

**Results:** We showed that CK2α’ haploinsufficiency in zQ175 mice ameliorated neuroinflammation, HTT aggregation, transcriptional alterations, excitatory synaptic transmission, and motor coordination deficits. RNA-seq analyses also revealed a connection between α-syn, a protein associated with PD, and the transcriptional perturbations mediated by CK2α’ in HD. We also found increased α-syn serine 129 phosphorylation (pS129-α-syn), a post-translational modification linked to α-synucleinopathy, in the nuclei of MSNs in zQ175 mice and in patients with HD. Levels of pS129-α-syn were ameliorated in zQ175 lacking one allele of CK2α’.

**Conclusions:** Our data demonstrated that CK2α’ contributes to transcriptional dysregulation of synaptic genes and neuroinflammation in zQ175 mice and its depletion improved several HD-like phenotypes in this mouse model. These effects were related to increased phosphorylation of S129-α-syn in the striatum of HD mice, suggesting that CK2α’ contributes to worsening HD by mediating synucleinopathy. Our study highlights a possible convergent mechanism of neurodegeneration between HD and PD and suggests targeting CK2α’ as a potential therapeutic strategy to ameliorate synaptic dysfunction in HD as well as other neurodegenerative diseases.

## Introduction

Huntington’s disease (HD) is a neurodegenerative disorder that manifests with progressive motor, cognitive, and psychiatric deficits for which there is no cure. HD is caused by a poly-glutamine (polyQ) expansion in exon 1 of the Huntingtin (*HTT*) gene. This mutation results in progressive misfolding and aggregation of mutant HTT protein (mtHTT) and preferentially affects GABAergic medium spiny neurons (MSNs) in the striatum (1–3). Transcriptional perturbations in synaptic genes and neuroinflammation are key processes that precede MSN death and motor symptom onset (4). However, our understanding of the interplay between these processes, mtHTT aggregation, and their contributions to MSN susceptibility in HD is still incomplete.

Protein kinase CK2 is at the crossroads between neuroinflammation, protein aggregation, and synaptic activity, and has recently emerged as a potential therapeutic target of neurodegeneration (5–7). CK2 is a highly conserved serine/threonine kinase composed of two regulatory beta (CK2β) subunits and two catalytic subunits, alpha (CK2α) and alpha prime (CK2α’) (8, 9). The two catalytic subunits share high structural homology, but they differ in their tissue distribution and their ability to phosphorylate different substrates (10, 11). Our previous work showed that CK2α’, but not CK2α, is induced in HD MSNs and contributes to the dysregulation of protein quality control systems and HTT aggregation in cells and mouse models of HD (12, 13). However, other studies conducted *in vitro* have suggested a protective role of CK2 in HD via HTT phosphorylation (14, 15), imposing the necessity to clarify the specific involvement of CK2α’ in HD pathogenesis and its potential as a therapeutic target in HD.

CK2 is involved in the phosphorylation and aggregation of other pathological proteins like microtubule associated protein tau (MAPT) and alpha-synuclein (α-syn), proteins involved in Alzheimer’s (AD) and Parkinson’s disease (PD) (16, 17). Phosphorylation of Tau and α-syn contribute to the activation of neuroinflammatory processes, transcriptional dysregulation, and synaptic deficits in AD and PD (18, 19). Alterations in these proteins have also been associated with HD pathology (20–22). In particular, increased levels of α-syn were observed in the plasma of patients with HD (23) and its deletion in R6/1 mice resulted in amelioration of motor deficits (20, 24). However, the mechanisms by which these proteins are altered in HD and the extent to which they contribute to HD pathophysiology are still unknown.

In this study, we characterized the role of CK2α’ in HD *in vivo* by using the heterozygous zQ175 HD mouse lacking one allele of CK2α’. We showed that CK2α’ haploinsufficiency decreased the levels of pro-inflammatory cytokines and improved astrocyte health, restored synaptic gene expression and excitatory synapse function, and improved motor behavior in zQ175 mice. These neuropathological and phenotypic changes correlated with alterations in α-syn serine 129 phosphorylation (pS129-α-syn) in the striatum, a post-translational modification involved in α-synucleinopathy, establishing a novel connection between CK2α’ function and synucleinopathy in HD. Collectively, our data demonstrated that CK2α’ plays a negative role in HD and indicates the therapeutic potential of modulating CK2α’ to achieve enhanced neuronal function and neuroprotection.

## Results

### Increased CK2α’ levels in the striatum of zQ175 mice parallel progressive HTT aggregation and NeuN depletion

Increased CK2 activity has been associated with detrimental effects in protein homeostasis and neuroinflammation in different neurodegenerative diseases, but its role in HD is still controversial (12, 14, 15). To determine whether CK2α’ plays a negative role during HD pathogenesis, we first evaluated the relationship between HTT aggregation, neuronal loss, and CK2α’ levels in the striatum over time for the heterozygous zQ175 mouse model at 3 (pre-symptomatic), 6 (early symptomatic), 12 (symptomatic), and 22 months (late-stage disease) of age (25, 26). We observed an age-dependent increase of HTT aggregates (EM48^+^ puncta) and fewer NeuN^+^ neurons (pan-neuronal marker) in the striatum of zQ175 mice (Fig. 1A-D**, S1A-C**). Increased HTT aggregates were also seen over time in the cortex of zQ175 mice, but they were delayed and significantly lower than in the striatum (**Fig. S1A, B**), as previously described (27). We confirmed that the depletion of NeuN^+^ cells correlated with decreased Ctip2^+^ neurons (MSN marker) (28)(**Fig. S1D-E**). However, we did not observe a significant difference in the total number of neurons, measured by cresyl violet (**Fig. S1F-H**), or in striatum volume (**Fig. S1I, J**), suggesting that changes in NeuN and Ctip2 reactivity may reflect transcriptional dysregulation and/or neuronal dysfunction rather than neuronal loss.

**Figure 1.**
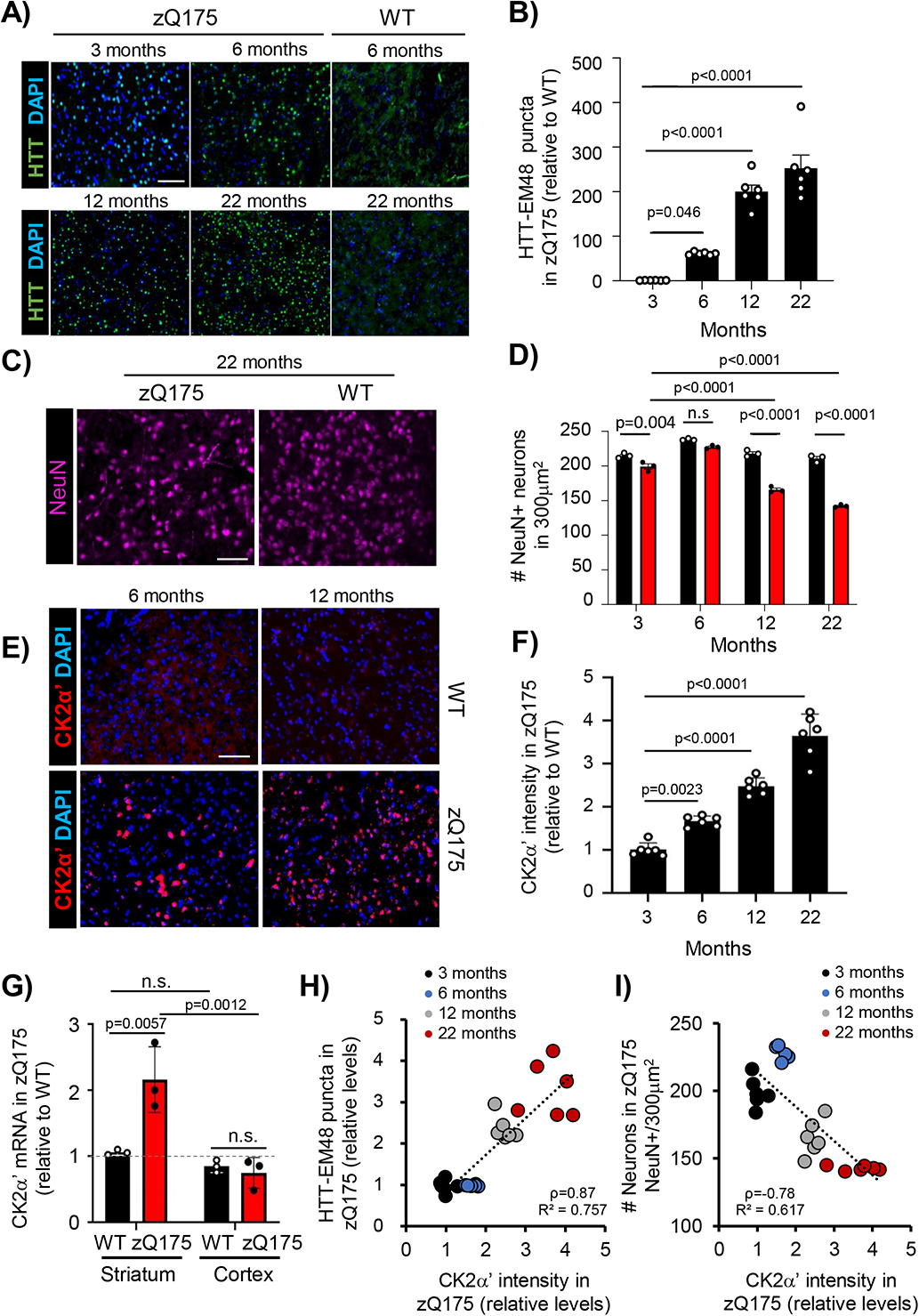
CK2α’ levels progressively increase in the striatum of zQ175 and correlate with HTT aggregation and NeuN depletion. **a**-**f**, Immunostaining and quantification of HTT puncta detected with anti-HTT EM48 antibody (*n*=6 mice/genotype) (**a**, **b**), NeuN^+^ cells (*n*=3 mice/genotype) (**c**, **d**) and CK2α’ levels (*n*=6 mice/genotype) (**e**, **f**) in zQ175 compared with WT mice at 3, 6, 12 and 22 months. **g**, CK2α’ mRNA levels analyzed by RT-qPCR in striatum and cortex of 6-month-old mice. Data was normalized to GAPDH and WT striatum (n=3 mice/genotype). **h**, Linear regression analysis between CK2α’ levels and HTT puncta, and ***i***, between CK2α’ levels and number of NeuN^+^ cells in zQ175 mice. The Pearson correlation coefficient (ρ) and *R^2^* are indicated. Data are mean ± SEM with significance determined by one-way ANOVA with Dunnett’s post-hoc test in **b**, mean ± SD with significance determined by one-way ANOVA Dunnett’s post-hoc test in **f** and two-way ANOVA withTukey’s post-hoc test in **d** and **g**. p-values <0.05 are indicated. n.s = not significant. Scale bar, 50 µm.

Due to the differences observed in the timing and level of HTT aggregation between striatum and cortex (**Fig. S1A, B**), we hypothesized that specific up-regulation of CK2α’ in the striatum contributes to the enhanced accumulation of HTT aggregates in the striatum. The levels of CK2α’ increased over time in zQ175 mice in the striatum but not in the cortex (Fig. 1E-G), coinciding with the timing of HTT aggregation and preceding robust NeuN depletion in the striatum. Regression analysis demonstrated that CK2α’ levels had a significant positive relationship with HTT aggregation (Pearson r(22)=0.87, p value<0.001) (Fig. 1H) and a significant negative relationship with the number of NeuN^+^ cells (Pearson r(22)=-0.78, p value<0.001) (Fig. 1I).

### Depletion of CK2α’ improves neuronal function and motor coordination

CK2 has been involved in the regulation of glutamate receptor trafficking via phosphorylation of receptor subunits as well as scaffolding proteins, suggesting a role of CK2 in neuronal signaling (29, 30). In addition, upregulation of the CK2α’ subunit in HD has been associated with alterations in MSN spine maturation and striatal synapse density in HD mice (12). Based on this evidence we decided to explore the functional extent of CK2α’ in HD by using a zQ175 mouse model lacking one allele of CK2α’ (zQ175:CK2α’^(+/-)^) (12) (Fig. 2A, B). We first assessed MSNs abundance and striatal synaptic proteins expression (**Fig. S2A-C**). CK2α’ haploinsufficiency in zQ175 mice did not alter the number of MSNs (Ctip2^+^ cells) or the mRNA levels of the MSN markers (Drd1 and Drd2), but increased the levels of synaptic proteins like the scaffold protein Dlg4 (PSD-95) and Ppp1rb1 (dopamine-and cAMP-regulated neuronal phosphoprotein DARPP-32), a key regulator of the electrophysiological responses in striatal neurons (32, 33) (**Fig. S2A-C**).

**Figure 2.**
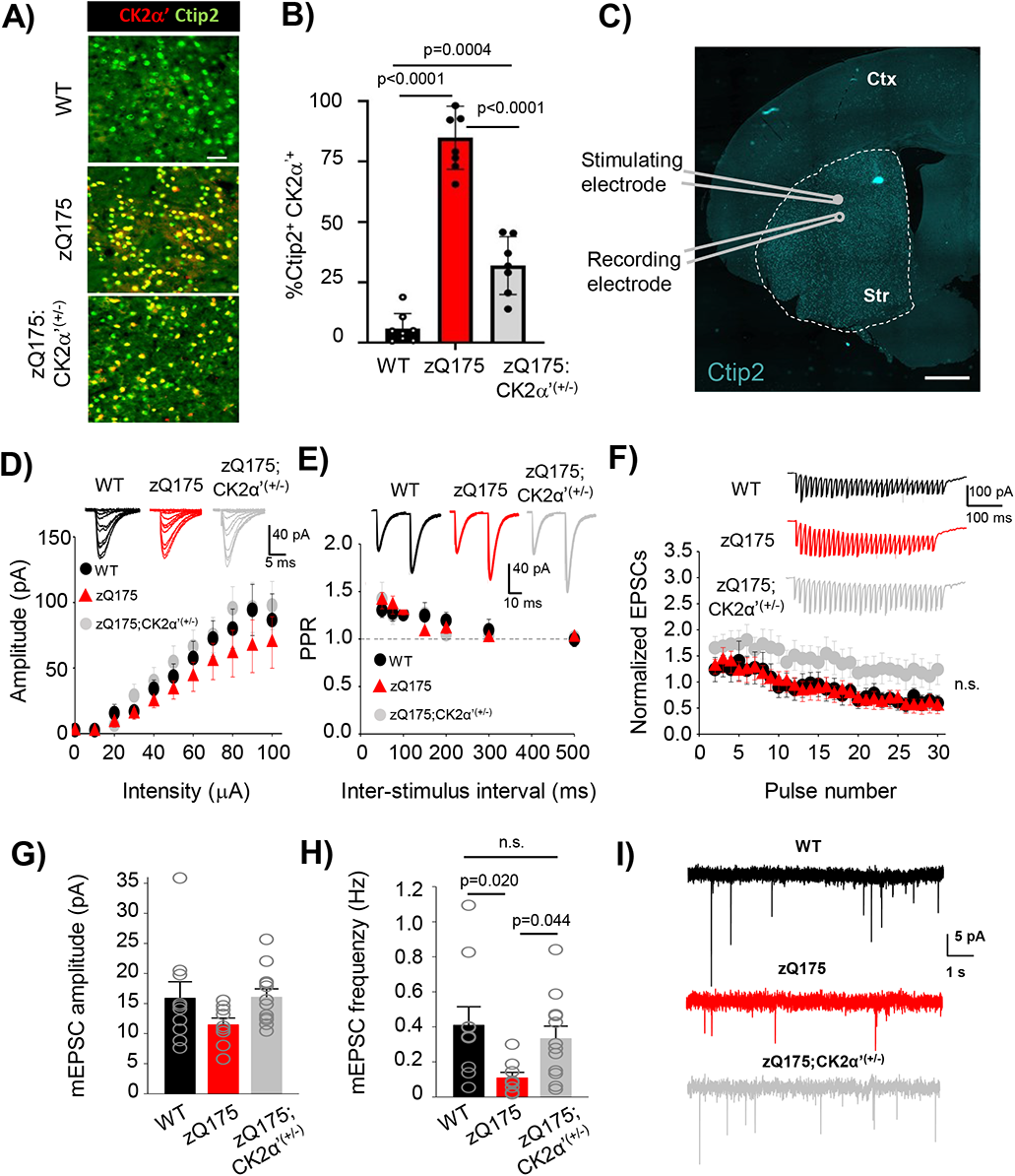
CK2α’ haploinsufficiency increased the frequency of AMPA-mediated miniature excitatory postsynaptic currents (mEPSC) in the dorsolateral striatum of zQ175 mice. **a**, **b**, Representative images show the labeling (**a**) and quantification (**b**) of CK2α’ in striatal MSNs immunostained for Ctip2, a specific MSN marker in WT, zQ175 and zQ175:CK2α’ ^(+/-)^ mice at 12 months of age (*n*=6 mice/genotype). Scale bar, 50 µm. **c**, Image shows whole-cell patch-clamp recording diagram in acute dorsolateral striatum slices, where Ctip2 labeled MSNs from 12-month-old mice. Scale bar 500 µm, Ctr: Cortex; Str: Striatum. **d**, Input–output curve (WT, n = 8; zQ175, n = 9; zQ175:CK2α’^(+/-)^ n = 13). Representative traces are shown in the top inset. **e**, Short-term potentiation measured via paired-pulse facilitation (WT, n = 8; zQ175, n = 9; zQ175:CK2α’^(+/-)^ n = 11). Representative traces of two consecutive stimuli delivered at 25 ms time intervals are shown in the top inset. **f**, Short-term depression analyzed through synaptic fatigue (WT, n = 7; zQ175, n = 9; zQ175:CK2α’^(+/-)^ n = 12). Representative traces are shown in the top inset. Values were analyzed using two-way ANOVA with Tukey’s post-hoc analysis. **g**, **h**, Spontaneous recordings of mini excitatory postsynaptic currents (mEPSCs). Amplitude (in pA; left panel) (**g**) and frequency (in Hz; right panel) (**h**) were analyzed (WT, n = 10; zQ175, n = 9; zQ175:CK2α’^(+/-)^ n = 12). **I**, Representative mEPSC traces. Values were analyzed using one-way ANOVA with Dunńs post-hoc analysis. P values <0.05 are indicated. Error bars represent mean ± SEM from at least 3 mice/genotype.

We then assessed the impact of CK2α’ depletion in AMPA-mediated excitatory transmission by conducting whole-cell patch clamp recordings from acute dorsolateral striatum coronal slices at 12 months (Fig. 2C). MSNs from all genotypes showed similar profiles in the analysis of basal synaptic transmission, including input/output curves, paired-pulse facilitation, and synaptic fatigue (Fig. 2D-F). We observed a trend towards increased normalized excitatory postsynaptic currents (EPSCs) in zQ175:CK2α’^(+/-)^ mice compared to the other two genotypes, but the data did not reach statistical significance (Fig. 2F). Spontaneous neurotransmitter release and synaptic activity via miniature EPSC (mEPSC) recordings showed that mEPSC amplitude, reflecting postsynaptic AMPA receptor function, was comparable among the 3 genotypes (Fig. 2G). However, mEPSC frequency, which reflects the probability of neurotransmitter release from presynaptic vesicles and also correlates with the number of synapses, was reduced in zQ175 mice (Fig. 2H, I), as previously reported (31), and rescued in zQ175:CK2α’^(+/-)^.

These data supported the role of CK2α’ in the dysregulation of striatal synaptic activity and excitability in HD mice. Glutamatergic synaptic transmission is often related to motor and cognitive function in HD mouse models (33, 34). We conducted a series of motor tests including accelerating rotarod and beam walk in WT, zQ175, and zQ175:CK2α’^(+/-)^ mice at 3, 6, and 12 months (Fig. 3). We also conducted cylinder and open field assessments on a different cohort at 12 months comparing zQ175 and zQ175:CK2α’^(+/-)^ (**Fig. S3**). We did not observe significant differences between WT and zQ175 or between zQ175 and zQ175:CK2α’^(+/-)^ at any tested age in the accelerating rotarod test (Fig. 3A-C), open field, or cylinder test (**Fig. S3**). However, when we evaluated fine motor coordination and whole-body balance in the beam test, we observed a significant increase in foot slips of zQ175 mice compared to WT at 3 months, but only with the most challenging beam (small round), indicating early subtle motor deficits (Fig. 3D). At 12 months, zQ175 mice showed increased foot slips in both the small round and small square beams compared to WT, highlighting a worsening motor deficit (Fig. 3F). zQ175:CK2α’^(+/-)^ mice showed a significant reduction in foot slips compared to zQ175 mice at all tested ages and no significant differences compared to WT.

**Figure 3.**
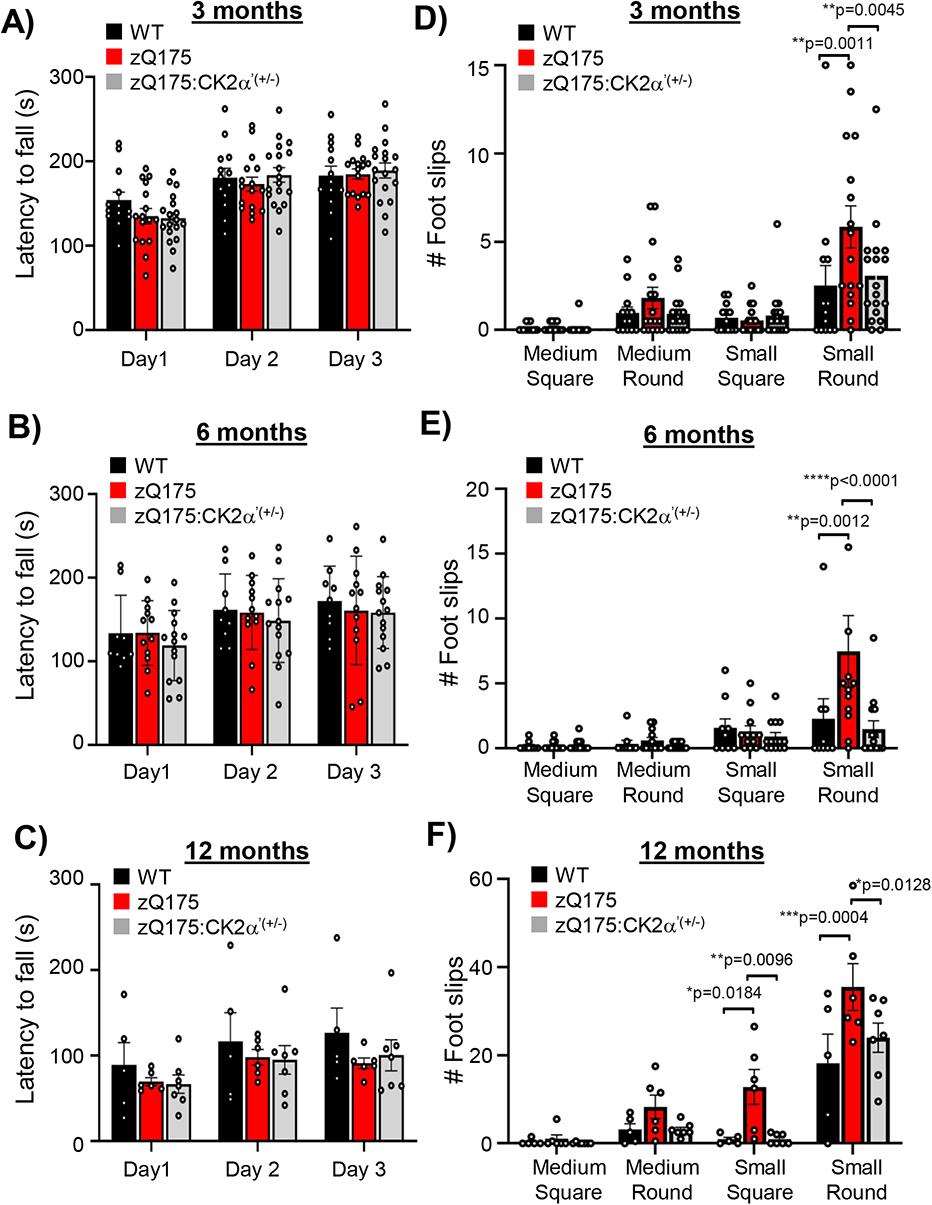
Genetic deletion of CK2α’ improved motor coordination in zQ175. **a-c**, Latency to fall off the rod (Rotarod test) for three consecutive days. **d-f**, Number of foot slips recorded while walking on four different types of beams with different degrees of difficulty from less to more challenging: medium-square, medium round, small-square and small-round (Beam test). Analyses were performed at 3, 6 and 12 months of age (n =16-18 mice/genotype in 3 months, n= 12-14 for 6 months and n = 5-6 for 12 months). Error bars denote mean ± SEM, values were analyzed by two-way ANOVA with *Sidak’s* post-hoc test. p-values <0.05 are indicated, n.s = not significant.

We also performed tests to evaluate associative learning (fear conditioning), spatial learning and memory (Barnes maze, BM), cognitive flexibility (BM reversal), and spatial working memory (Y radial arm maze) by comparing zQ175 and zQ175:CK2α’^(+/-)^ mice at 12 months of age, but no significant differences were observed between the two groups (**Fig. S4**). This observation suggests that the positive effects of CK2α’ depletion on motor behavior may not additionally translate to improved cognitive functions.

### CK2α’ depletion rescued transcriptional dysregulation of genes involved in glutamatergic signaling

We sought to determine whether depletion of CK2α’ levels had any influence in the overall transcriptional dysregulation characteristic of HD and whether those changes could be associated with the functional improvement observed in zQ175:CK2α’^(+/-)^ mice. We performed RNA-seq in the striatum of 12-14 month old mice, followed by Weighted Gene Co-Expression Network Analysis (WGCNA) to investigate which molecular pathways are affected by CK2α’ using n=5 mice/genotype for WT, zQ175, and zQ175:CK2α’^(+/-)^ and n=3 mice for CK2α’^(+/-)^. We found that the mouse transcriptome could be clustered into 20 gene co-expression modules (**Fig. S5, Table S1**). Nine modules showed a significant difference in eigengene expression between zQ175 and WT in a Kruskal-Wallis test (p value < 0.05) (**Table S2, Fig. S6A**) and two modules (Greenyellow: 255 genes, and Red: 639 genes) were significantly different between zQ175 and zQ175:CK2α’^(+/-)^ mice (p value < 0.05) (Fig. 4A, B, **Table S2**). Cook’s distance (DESeq2) analyses revealed that these differences were not due to the presence of outliers in our data set (**Fig. S6B**). We focused our analyses on the Greenyellow module due to its higher significance. Ingenuity pathway analysis (IPA) indicated that the five most significant pathways in the Greenyellow module were signaling pathways for synaptogenesis (p-value 1.68E-06), Ephrin A (p-value 7.84E-05), glutamate receptor (p-value 1.98E-04), axonal guidance (p-value 7.13E-04), and G-protein coupled receptor (GPCR) (p-value 1.14E-03) (Fig. 4C), all of which are pathways previously shown to be dysregulated in HD (35). IPA in the Red module also revealed synaptic signaling related pathways among their five most significant pathways (**Fig. S6C**). Additional Gene Ontology (GO) annotation of cellular components of the Greenyellow indicated that genes were enriched in synaptic components (Fig. 4D).

**Figure 4.**
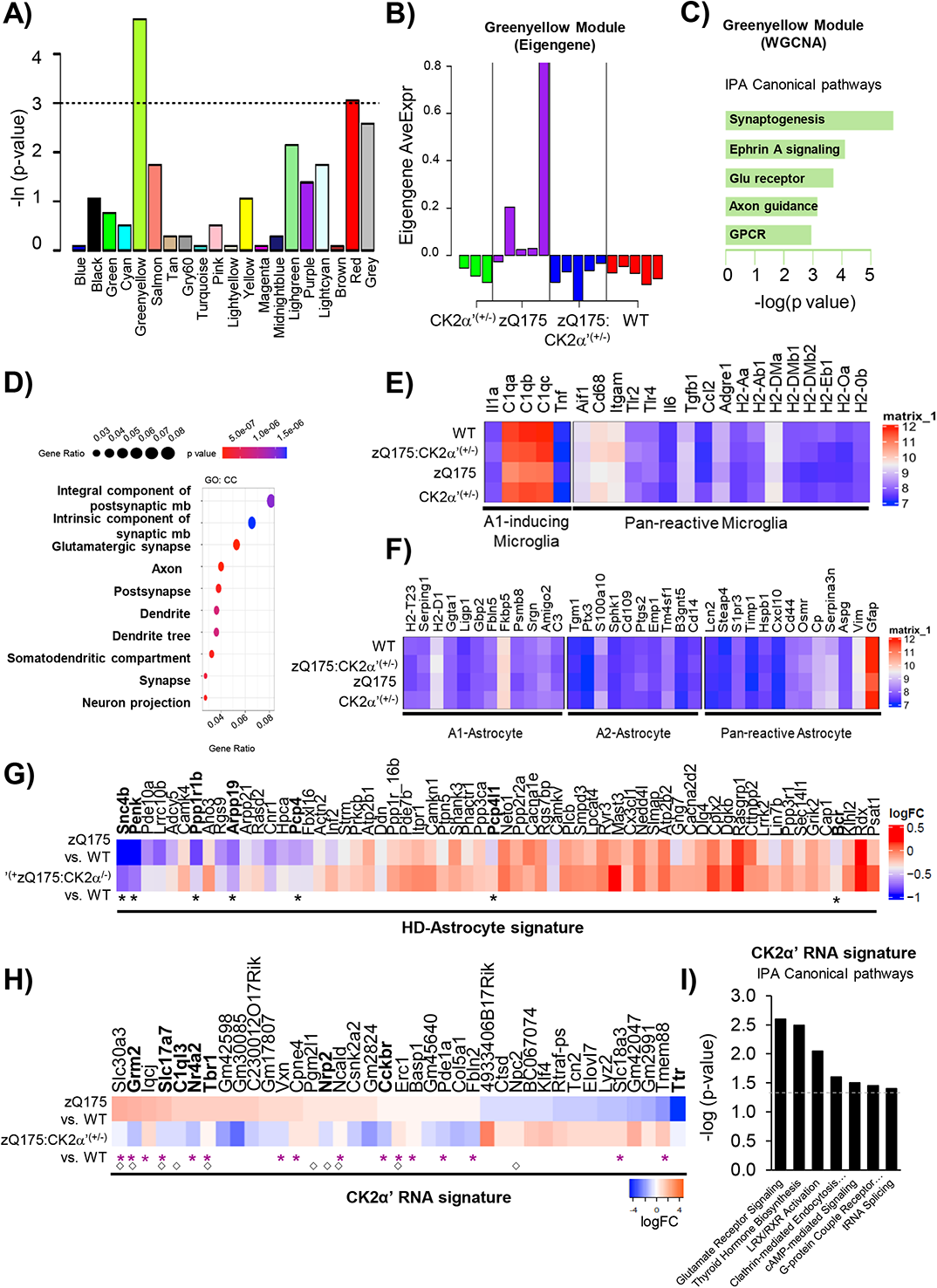
Depletion of CK2α’ restored the expression of synaptic genes associated with α-syn-dependent regulation in the striatum of symptomatic zQ175 mice. **a**, Kruskal-Wallis test of module expressions between zQ175 (HD) mice and zQ175:CK2α’^(+/-)^ mice. The y-axis is the negative log transformed p-values. **b**, Expressions of module “Greenyellow” in each mouse sample. **c**, IPA canonical pathway analysis, **d**, enrichment analysis of GO terms in CC (cellular component). ***e-f***, Gene expression for microglia marker genes; A1-inducing and pan-reactive microglia genes (40) (**e**) and astrocyte markers representative of A1, A2 and pan-reactive astrocytes genes (40) (**f**) in WT, zQ175, CK2α’^(+/-)^ and zQ175:CK2α’^(+/-)^ mice. **g**, **h**, Mean log2 fold change between zQ175 and zQ175:CK2α’^(+/-)^ mice compared to WT for genes representative of the HD-astrocyte molecular signature (41). (**g**) and the CK2α’-mediated RNA signature (**h**). Purple asterisk (*) indicates synaptic function, (◊) indicates genes present in the Greenyellow module. **i**, IPA canonical pathway analysis for the CK2α’-mediated RNA signature.

Connectivity analyses (**Fig. S6D**) revealed that the two most connected genes within the hub were Slit1 (Slit Guidance Ligand 1), associated with “poor” behavior and a worse prognosis in the R6/1 mouse mode (36), and Ncald (Neurocalcin delta), which regulates multiple endocytosis-dependent neuronal functions and is situated on a locus that has been associated with earlier clinical onset of HD (37, 38). Differential Gene Expression (DGE) between WT and zQ175 mice confirmed a large transcriptional dysregulation (n=885 genes, Q<0.1) (**Fig. S6E, F, Table S3**), as previously reported (35) while the DGE between zQ175:CK2α’^(+/-)^ and WT mice only reported 123 genes (**Fig. S6G, H**). R package variance Partition confirmed that these expression changes were driven only by genotype and not by differences in sex distribution among our groups (**Fig. S6I**).

CK2 has been previously associated with neuroinflammatory processes (6, 39), which was supported by the amelioration in the levels of inflammatory cytokines upon reduction of CK2α’ in both HD cells and mice (**Fig. S7A-D**). Therefore, we examined our data set for microglial and astrocytic inflammatory RNA signatures (40) but did not observe significant changes in the expression of these gene signatures across genotypes (Fig. 4E, F**, Table S4, S5**). Immunoblotting analyses of the microglial marker Iba1 (Ionized calcium binding adaptor molecule), considered a reactive marker of microgliosis, indicated an increase in total Iba1 protein levels between WT and the HD groups but immunohistological analyses of Iba1 showed no differences in the number or area size of Iba1^+^ cells across all genotypes (**Fig. S7E-H**), in line with the results obtained by RNA-seq (Fig. 4E). The discrepancy between changes in protein levels of inflammatory cytokines and the absence of an inflammatory transcriptional signature could be related to post-translational events potentially regulated by CK2α’. In addition, no changes in the RNA signature characteristic of reactive neurotoxic A1 astrocytes were seen across genotypes in our data set (Fig. 4F). The absence of robust microglial and astrocytic inflammatory RNA signatures in zQ175 and other HD models has previously been demonstrated (41). However, we recapitulated some transcriptional changes for the so called ‘*HD-associated astrocyte molecular signature*’ (41, 42) (Fig. 4G). Among all genes, zQ175 mice showed a significant decreased expression of Snc4b, Penk, Pppp1r1b, Arpp19, Pcp4, Pcp41 and Bcr, compared to WT mice, indicative of astrocytic dysfunction (Fig. 4G). Notably, these changes were ameliorated when comparing zQ175:CK2α’^(+/-)^ and WT mice, suggesting diminished astrocytic pathology upon reduction of CK2α’ levels. These results were supported by the amelioration of astrogliosis (**Fig. S8A, B**) and the reduction in the astroglia marker myo-inositol, measured by *in vivo* proton magnetic resonance spectroscopy (^1^H-MRS) (25, 43, 44), when comparing zQ175:CK2α’^(+/-)^ and zQ175 mice (**Fig. S8C-F**).

The DGEs analyzed between zQ175 and zQ175:CK2α’^(+/-)^ revealed 39 specific and significant genes (FDR<0.1) (CK2α’ RNA signature) (Fig. 4H**, Table S6**), which included Csnk2a2 (CK2α’ gene) as a positive control. Three genes (Ncald, Nrp2, and Slc30a3) were also among the 15% most highly connected members of the Greenyellow module (**Fig. S6I**). At least 40% of the DGEs (n=16) were related to synaptic functions (**Table S6**). IPA on the 39 genes showed that the most significant canonical pathway was for glutamate receptor signaling (p-value 2.59E-03) (Fig. 4I), confirming the contribution of CK2α’ to the dysregulation of genes related to excitatory synaptic transmission in HD. However, the available information for the regulation of the 39 gene set did not provide a direct connection between any of these hits and CK2α, therefore suggesting additional regulators implicated in the CK2α’-mediated RNA signature.

### α-syn participates in CK2α’-mediated synaptic gene dysregulation

When looking at the most significant upstream regulators identified by IPA of both the Greenyellow module and the 39 gene set identified by DGE, we found SNCA (α-syn) (p-value 9.10E-11 and 1.03E-07, respectively). α-syn regulates multiple processes including synaptic vesicle trafficking, neurotransmitter release and transcription (45, 46), and has been previously connected with CK2 (17, 47). IPA connected α-syn with some of the most differentially dysregulated genes by CK2α’ including Ttr (Transthyretin), Grm2 (Glutamate Metabotropic Receptor 2), Slc17a7 (Solute Carrier Family 17 Member 7; alias VGlut1), C1ql3 (Complement Component 1, Q Subcomponent-Like), Cckbr (cholecystokinin B receptor), Nrp2 (Neuropilin 2), and the transcription factors Tbr1 (T-Box Brain Transcription Factor 1) and Nr4a2 (Nuclear Receptor Subfamily 4 Group A Member 2; alias Nurr1) (Fig. 5A).

**Figure 5.**
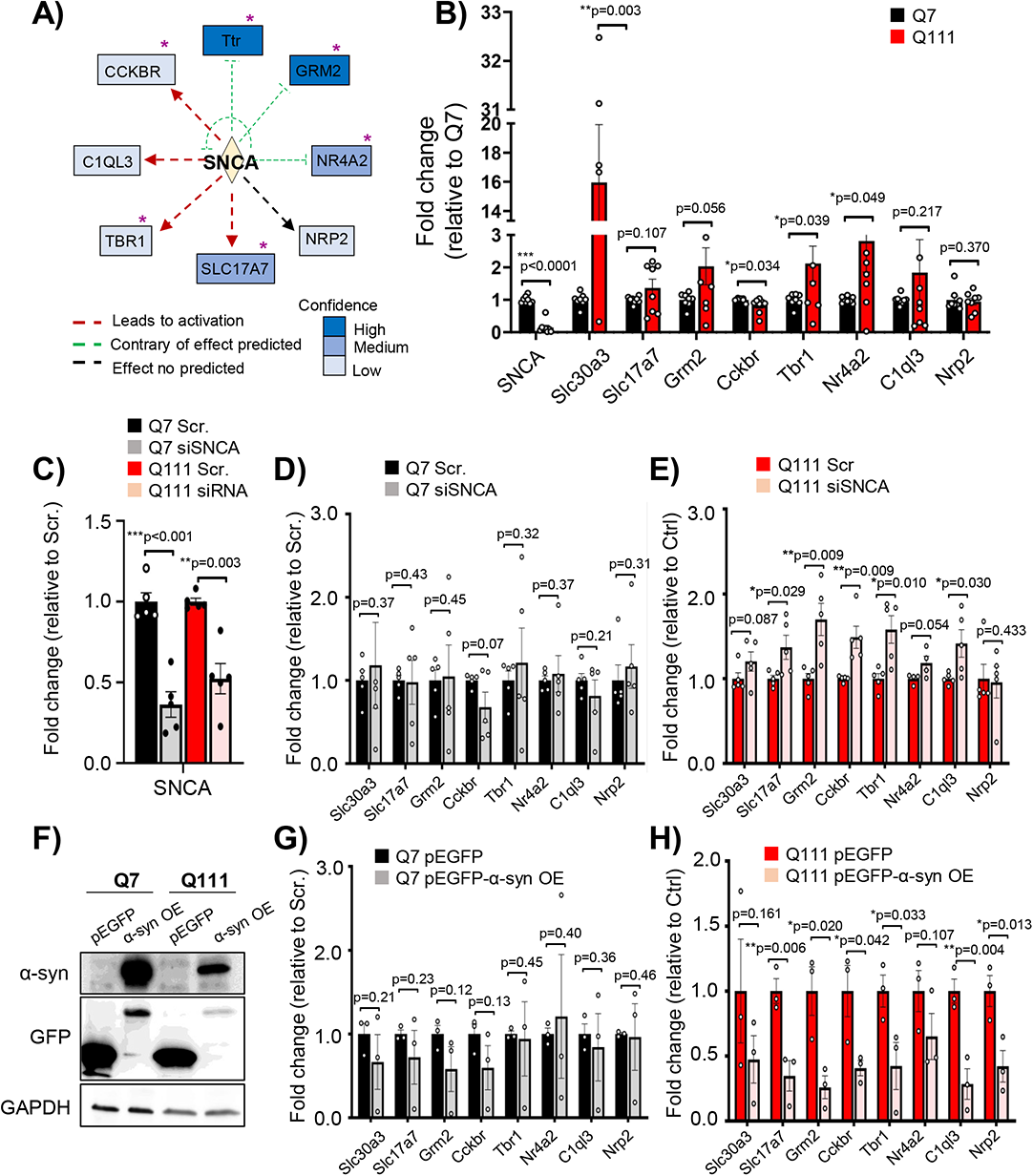
SNCA regulates the expression of genes identified in the CK2α’ mediated RNA signature. **a,** SNCA and DGEs connection by IPA network analysis. Purple asterisks denote synaptic function. **b**, RT-qPCR in Q7 and Q111 cells (*n* = 6-8 experiments). **c-e,** siRNA knockdown of SNCA (siSNCA) for 24 h in ST*Hdh Q7 (**d**)* and Q111 cells (**e**), and RT-qPCR for SNCA (**c**) and SNCA putative gene targets (**d, e**). Data were normalized with GAPDH and relativized to non-targeting control siRNA-treated cells (Scr.), (*n* = 5 experiments). Four data points fell in the axis break for Slc30a3. **f**, Immunoblotting for α-syn, GFP and GAPDH in Q7 and Q111 cells after plasmid transfections. Q7 and Q111 cells were transfected with pEGFP (control) or α-syn-GFP overexpression (OE) and harvested 24 h after transfection. **g**, **h**, RT-qPCR for Q7 (**g**) and Q111 cells (**h**) transfected with control or a-syn-GFP. Data were normalized with GAPDH and relativized to non-targeting control pEGFP-treated cells, (*n* = 3 experiments). Data are mean ± SEM with significance determined by Welch’s t-test.

To determine the extent to which α-syn participates in the regulation of genes identified in the CK2α’-mediated RNA signature by IPA, we silenced or overexpressed SNCA in the murine striatal cell models Q7 (control) and Q111 (HD) cells. We first validated that Q111 cells presented similar gene expression alterations to those observed in zQ175 mice for the putative SNCA targets when compared to Q7 cells (Fig. 5B). Slc30a3 was included as a non-SNCA target control. Ttr expression was not detected in either Q7 or Q111. RT-qPCR showed a significant increase in Slc30a3, Slc17a7, Grm2, Cckbr, Tbr1 and Nr4a2 in Q111 compared to Q7 as observed in zQ175 mice when compared with WT. Interestingly, SNCA transcripts in Q111 cells were significantly lower compared to Q7 (Fig. 5B). Silencing SNCA in Q111 cells significantly increased the expression of several putative SNCA targets; Slc17a7, Grm2, Cckbr, Tbr and C1ql3, but not the non-SNCA targeted control gene Slc30a3. No significant effects on Q7 cells were observed (Fig. 5C-E). On the contrary, α-syn overexpression (OE) in Q111 cells had opposite effects on the same SNCA target genes with no effect on Q7 cells. We also conducted analyses in R6/1 and R6/1:SNCA^KO^ mice compared to WT (**Fig. S9A, B**) (20). Although R6/1 mice did not show a similar transcriptional alteration for the SNCA target genes to that observed in zQ175, possibly due disease severity differences between these two mouse models, we observed significantly decreased SNCA transcripts in R6/1 mice compared to WT, as observed in Q111 compared to Q7 cells (**Fig. S9B**). We also observed that SNCA^KO^ significantly altered the expression of Grm2 in the R6/1 background but not in the WT background (**Fig. S9B**). Altogether, the effects mediated by SNCA manipulations suggested that transcriptional alterations of some synaptic genes in HD could be mediated by α-syn dysregulation.

### Striatal synucleinopathy is found in zQ175 mice and is reduced by CK2α’ depletion

We next explored whether CK2α’ was involved in the regulation of α-syn in HD. We observed the total amount of α-syn was similar between WT and zQ175 (Fig. 6A-C) mice. zQ175:CK2α’^(+/-)^ mice showed a trend towards increased α-syn, but did not reach statistical significance (Fig. 6B, C). Nuclear and cytoplasmic fractionation confirmed the presence of α-syn in nuclear fractions from striatum samples (45, 48), and showed a modest but significant increase in nuclear α-syn in zQ175 mice (Fig. 6D, E). IF analyses for α-syn and HTT (EM48) also confirmed the colocalization between these two proteins, as previously shown in R6/1 mice (20)(Fig. 6F, G). To determine if there was a difference in the number and distribution of co-localized α-syn/HTT, we first analyzed the number of EM48^+^ puncta in both the nucleus and cytoplasm between zQ175 and zQ175:CK2α’^(+/-)^ mice. Cytoplasmic HTT aggregates were reduced in zQ175:CK2α’^(+/-)^ compared to zQ175 mice, consistent with previous studies (12), although no significant differences were observed in the number of nuclear HTT aggregates (Fig. 6H, I). Despite the decrease in cytoplasmic HTT aggregates in zQ175:CK2α’^(+/-)^ mice, no significant differences were observed in the number of nuclear and/or cytoplasmic α-syn/HTT colocalized puncta between zQ175 and zQ175:CK2α’^(+/-)^ mice (Fig. 6J).

**Figure 6.**
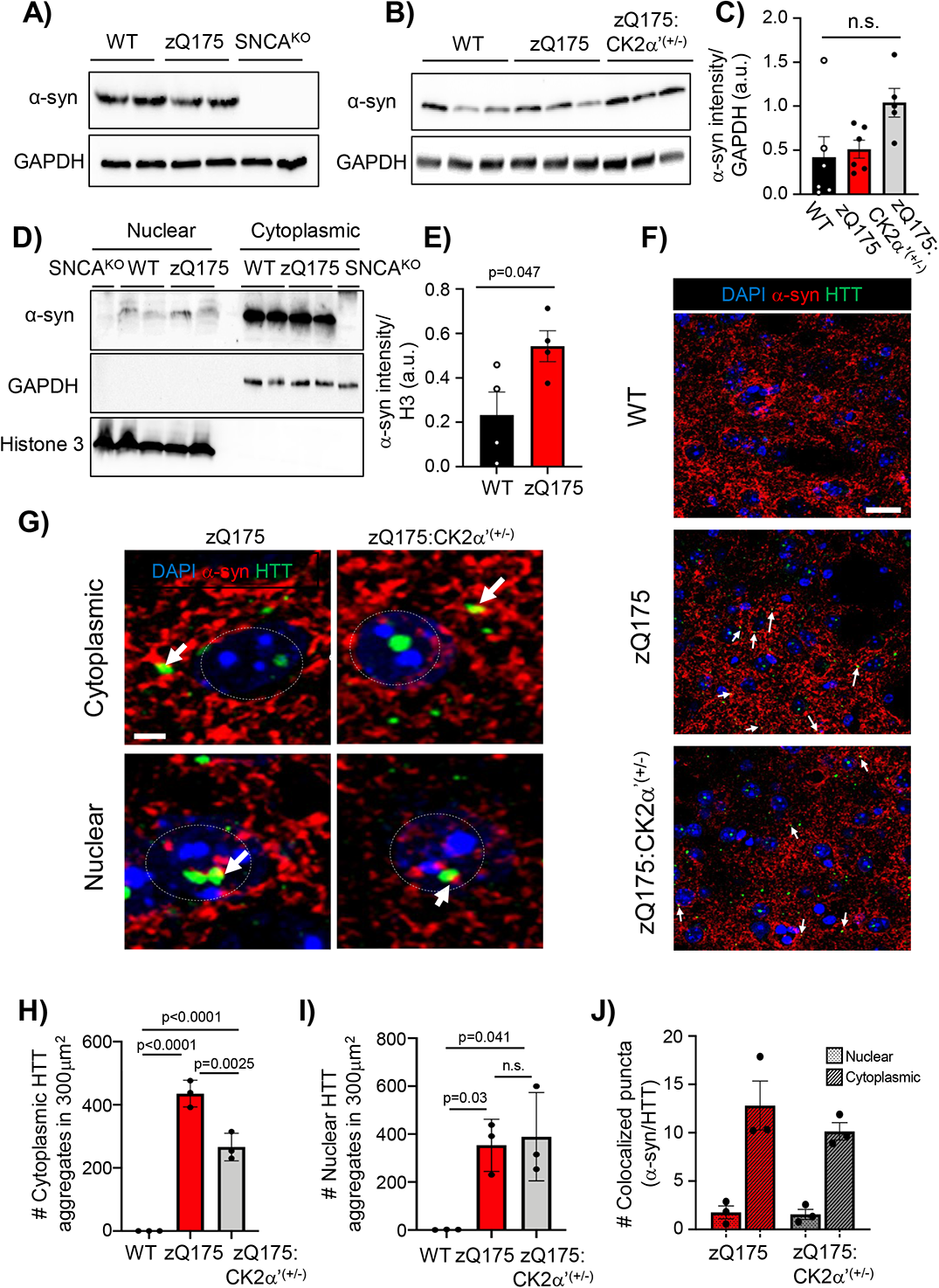
α-syn differentially accumulates in the nucleus of symptomatic zQ175 mice and colocalized with mtHTT. **a**, α-syn (4D6 antibody) IB in the striatum of WT, zQ175 and SNCA^KO^ and **b** in WT, zQ175 and zQ175:CK2α’^(+/-)^ mice at 12 months old. GAPDH used as loading control. **c**, α-syn protein levels analyzed by Image J from IB analyses (n= 5-6 mice/genotype). **d**, Nuclear/cytoplasmic fractionation of striatum samples from 12-month-old WT, zQ175 and SNCA^KO^ mice. **e**, Quantification of nuclear α-syn from images in D (n=4 mice/genotype). **f**, α-syn and HTT (EM48 antibody) IF images of dorsal striatum sections from 12 month old WT, zQ175 and zQ175:CK2α’^(+/-)^ (n=3 mice/genotype). White arrows indicate α-syn/HTT colocalization. Scale bar, 10 μm. **g**, Magnification of images from F. Scale bar, 2 μm. Grey circles represent nuclei. **h** Number of cytoplasmic and ***i*** nuclear EM48^+^ puncta. A total of three images per brain section and three brain sections per genotype were analyzed (n=27 images, n=3 mice/ genotype). **j**, Number of colocalized α-syn and EM48^+^ puncta calculated using Image J Puncta analysis plugin (n=3 mice/genotype). Error bars denote mean ± SEM, values were analyzed by Student’s t-test.

We then evaluated whether pS129-α-syn, a marker of synucleinopathy (49, 50), was altered in HD and whether CK2α’ could influence its levels. We observed that the levels of pS129-α-syn increased in the striatum of zQ175 mice at 12 months compared to WT (tested with 3 different pS129-α-syn antibodies: 81A, EP1536Y and D1R1R), and in the striatum of patients with HD (Fig. 7A-E**, Fig. S10A**), indicating signs of synucleinopathy. The levels of pS129-α-syn were significantly reduced in zQ175:CK2α’^(+/-)^ mice compared to zQ175, while no significant differences were observed with WT mice (Fig. 7D, E, **Fig. S10A-C**). pS129-α-syn was detected in both the cytoplasm and the nucleus of zQ175 striatal cells, while no nuclear presence was detected in zQ175:CK2α’^(+/-)^ mice (Fig. 7F, G). In addition, we observed that pS129-α-syn colocalized with both cytoplasmic and nuclear HTT puncta in zQ175 mice, while only cytoplasmic colocalization was observed in zQ175:CK2α’^(+/-)^ mice (Fig. 7F, G**, Suppl. Video 1**).

**Figure 7.**
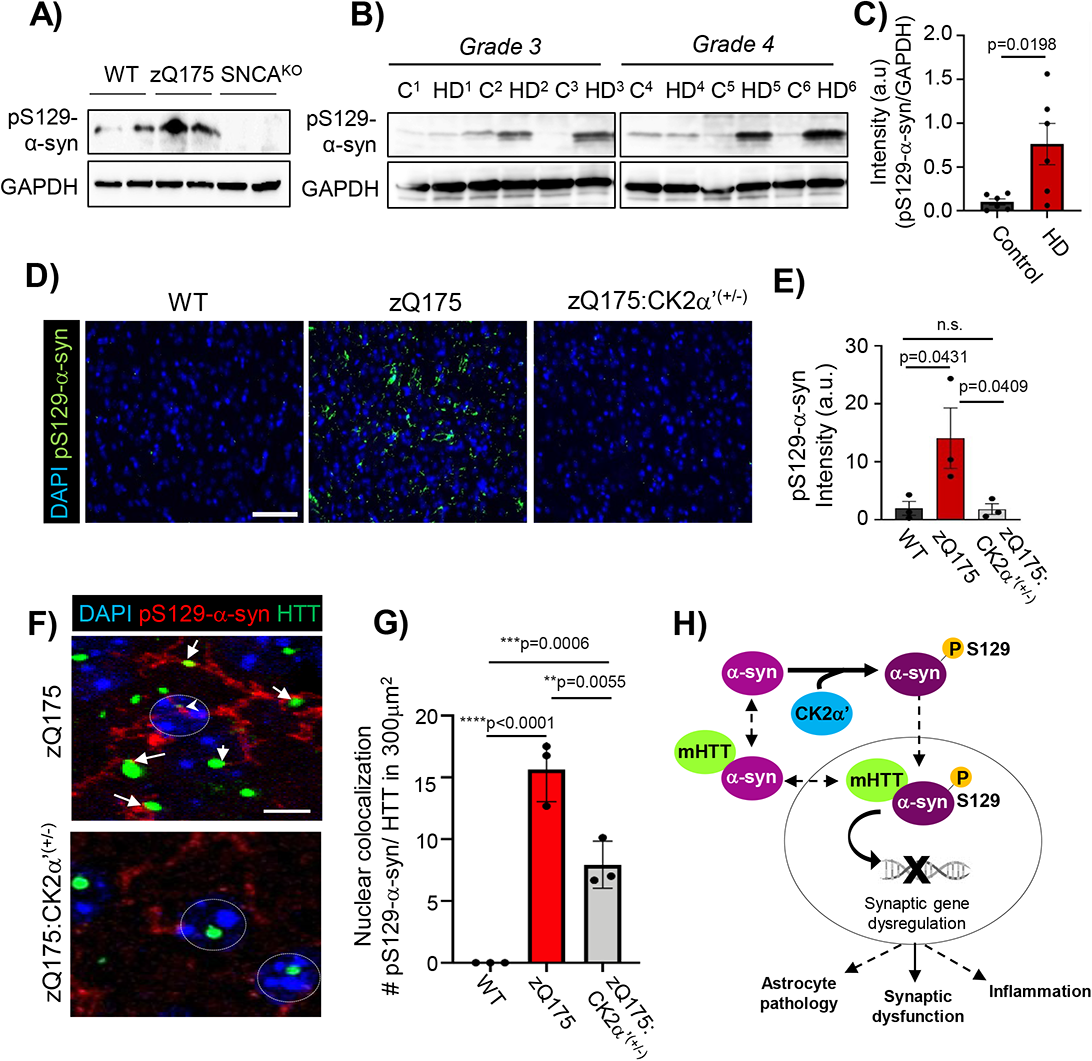
CK2α’ regulates phosphorylation of S129-α-syn and nuclear accumulation in symptomatic zQ175 mice. **a**, pS129-α-syn (EP1536Y antibody) IB in the striatum of 12-month-old WT, zQ175 and SNCA^KO^ (n=4 mice/genotype). **b**, pS129-α-syn (81A antibody) IB in the striatum of patients with HD (Vonsattel grade 3 and 4, Harvard Brain Tissue Resource Center) compared to age and sex matched controls. GAPDH is used as loading control. ***c***, pS129-α-syn protein levels (combined grades 3 and 4) analyzed by Image J from images in **b**. **d**, pS129-α-syn IF (81A antibody) in the dorsal striatum of 12-month-old WT, zQ175 and zQ175:CK2α’^(+/-)^ (n=3 mice/genotype), Scale bar, 20 μm. **e**, pS129-α-syn fluorescence signal was calculated using Image J from images in D (n=3 mice/genotype, n=27 images per mouse). **f**, Magnification of images in D, pS129-α-syn and EM48 colocalization in zQ175 and zQ175:CK2α’^(+/-)^ (n=3 mice/group). **g**, Quantification of pS129-α-syn and EM48 colocalized puncta using Image J puncta plug in. All data are mean ± SEM. Statistical analyses were conducted by one-way ANOVA. **h**, Working model for the role of CK2α’ in the regulation of pS129-α-syn and HD-like phenotype.

## Discussion

Increased protein kinase CK2 activity has been associated with detrimental effects in protein homeostasis and neuroinflammation in different neurodegenerative diseases, including AD and PD (5). However, the role of CK2 in HD remained unclear. We previously showed that CK2α’ is induced in cell and mouse models of HD, in human iPSC-MSN like cells derived from patients with HD and in postmortem striatum from patients with HD (12). Increased CK2 was also reported in polyQ-HTT expressing cells and in the YAC128 HD mouse model (14). CK2α’ genetic knockdown in different HD cell models resulted in decreased HTT aggregation and increased cell viability (12) but other studies using CK2 inhibitors resulted in opposite results and suggested CK2 had a protective role in HD (14, 15). These contradictory data claimed a more in-depth characterization of the role of CK2 in HD. Here we have demonstrated the adverse effects of CK2α’ catalytic subunit on several HD-related phenotypes including transcriptional dysregulation, neuroinflammation, protein aggregation, neuronal function, and motor coordination in the zQ175 HD mouse model and consolidated the detrimental contribution of CK2α’ to HD pathogenesis. We found CK2α’ contribution is mediated, at least in part, by the ability of CK2α’ to influence α-syn phosphorylation, striatal synucleinopathy and synaptic gene dysregulation (Fig. 7H).

CK2 has been widely associated with the activation of neuroinflammatory processes (39, 51, 52). One of the proposed mechanisms is the participation of CK2 in the phosphorylation of components of the IKK (IκB kinase)/NFκB pathway that results in the production of proinflammatory cytokines (52). Studies in AD showed that CK2 is involved in the inflammatory response that occurs in astrocytes and demonstrated that pharmacological inhibition of CK2 reduced pro-inflammatory cytokine secretion by human astrocytes (6). Although the role of CK2 in inflammation is mostly attributed to the CK2α subunit (39), we showed that CK2α’ haploinsufficiency reduced the levels of several proinflammatory cytokines and diminished astrogliosis in the striatum of zQ175 mice. The benefits of reducing neuroinflammation in HD have been shown in R6/2 mice intracranially injected with a TNF-α inhibitor, which resulted in improved motor function (53). Similarly, CK2α’ haploinsufficiency in zQ175 resulted in improved motor coordination.

Despite the beneficial effects on neuroinflammation and motor behavior observed by reducing the levels of CK2α’ in zQ175 mice, our transcriptomic analyses did not reveal a neuroinflammatory transcriptional response (40) in either zQ175 or zQ175:CK2α’^(+/-)^ mice. This was also supported by the absence of a robust microgliosis phenotype in these mice. While the disconnection between changes in protein levels of inflammatory cytokines and the absence of an inflammatory transcriptional signature is intriguing, similar results have previously been shown in zQ175 and other HD mouse models like the R6/2 (41). Transcriptomic studies in these mice have instead revealed a transcriptional signature related to astrocytic dysfunction (41). Our own RNA-seq data confirmed this HD astrocyte molecular signature manifested by the downregulation of several astrocyte-related genes including Snc4b, Penk, Ppp1r1b and Arpp19. We found that changes in CK2α’ levels influenced the expression of some of those genes associated with astrocytic dysfunction, suggesting a role of CK2α’ in HD-related astrocyte pathology.

Increased pro-inflammatory cytokines can alter synaptic strength as well as glutamatergic transmission and are also associated with structural and functional disruption of synaptic compartments (54). Similarly, astrocyte dysfunction in HD contributes to reduced striatal glutamatergic transmission and spine density, ultimately decreasing MSN excitability (55, 56). We showed that CK2α’ depletion increased the expression of synaptic proteins (PSD-95 and Darpp-32) and improved AMPA-mediated synaptic transmission. It is possible that these effects are mediated by an improved astrocyte health influenced by the reduction of CK2α’. In support of this hypothesis, conditional deletion of mtHTT in astrocytes of BACHD mice improved astrocyte health and rescued the expression of synaptic proteins like PSD-95 and improved striatal synaptic activity (55). However, whether decreased astrocytic pathology contributes to improved neuronal activity in zQ175:CK2α’^(+/-)^ mice is yet to be determined.

In contrast to CK2α, which is an essential protein with hundreds of targets, CK2α’ has very few identified substrates (10, 57). Based on pharmacological studies, it was proposed that CK2 protected cells by mediating Ser13 and Ser16 HTT phosphorylation and decreasing HTT toxicity (14, 15, 58), but genetic evidence for the direct role of CK2 (either CK2α or CK2α’ subunits) has never been shown. Previously used CK2 inhibitors were characterized by their high toxicity and poor selectivity with capacity to potently inhibit other kinases (59, 60). This could explain indirect effects on HTT phosphorylation and decreased cell viability when used in HD cells (14, 15). In addition, the N-terminal sequence of HTT (KAFE**<u>S</u>_13_**LK**<u>S</u>_16_**FQQQ) lacks the CK2 consensus sequence (SxxE/D) (10, 57). A recent kinase screening has revealed that other kinases like TBK1 are most likely to be involved in HTT phosphorylation than CK2 (61). Although we cannot rule out potential indirect effects of CK2α’ on HTT phosphorylation in zQ175 mice, we showed that genetic reduction of CK2α’ levels in both HD cells (12) and mouse models decreased HTT aggregation and toxicity, which is contrary to what would be expected if CK2α’ participates in HTT phosphorylation.

On the other hand, we previously showed CK2α’ directly phosphorylates the stress protective transcription factor HSF1, which regulates protein homeostasis (13), signaling HSF1 for proteasomal degradation and influencing chaperone expression in HD (12). Our RNA-seq analysis validated the increased expression of chaperones like Hsp70 and Hsp25 in zQ175:CK2α’^(+/-)^ mice, consistent with previous findings (12). However, WGCNA and DGE did not reveal global changes in transcriptional pathways associated with protein quality control networks in zQ175:CK2α’^(+/-)^ mice, but instead showed a unique CK2α’-mediated RNA signature related to synaptogenesis and glutamate receptor signaling. This data correlated with the improved frequency of striatal mEPSCs observed by reducing CK2α’ levels and supports previous findings showing increased MSNs maturation and striatal synapse density in zQ175:CK2α’^(+/-)^ mice (12, 62). α-syn was shown by IPA as one of the top putative upstream regulators of the CK2α’- mediated transcriptional changes. Although α-syn is not a transcription factor, several reports showed α-syn modulates transcription by either regulating the expression of transcription factors like Nurr1 (45, 46), which is differentially expressed between zQ175 and zQ175:CK2α’^(+/-)^, or by inducing epigenetic modifications in the DNA (63). Interestingly, mice expressing human α-syn selectively altered glutamate receptor signaling genes at both the epigenetic and transcriptional level (63), which supports the hypothesis that CK2α’-mediated alterations in glutamatergic signaling could be α-syn dependent.

α-Syn participates in HD pathogenesis since α-syn KO mice decreased mtHTT aggregation and attenuated body weight loss and motor symptoms in R6/1 mice (20, 64), although its specific mechanism of action in HD was not stablished. Aggregation of α-syn and consequent synucleinopathy in PD were linked to CK2-dependent phosphorylation of S129-α-syn (YEMP**S_129_**EEG), although this site is also the target of other protein kinases (47, 65). Here, we showed that pS129-α-syn levels are increased in the striatum of symptomatic HD mice and patients with HD as well as increased pS129-α-syn localization in the MSN nucleus. pS129-α-syn was decreased when reducing the levels of CK2α’. Increased pS129-α-syn in cortical neurons of aged mice has been correlated with the dysregulation of vesicular glutamate transporter Slc17a7 (66), which is also seen in zQ175 mice. Considering all the evidence, it is reasonable to hypothesize that CK2α’-mediated increase of pS129-α-syn in the brains of zQ175 mice could contribute to glutamate signaling dysregulation by altering (directly or indirectly) the expression of genes related to those processes and ultimately affecting several HD-like phenotypes. However, we cannot disregard the possibility that the effects mediated by CK2α’ depletion could be additionally influenced by other CK2α’ substrates. Further experiments will be necessary to decipher the mechanism by which CK2α’-mediated α-synucleinopathy contributes to HD and to tease apart the differential contribution of HTT aggregation and α-syn pathology to the symptomatology, onset, and progression of HD.

## Materials and Methods

See SI Appendix for complete methods.

### Cell lines

Mammalian cell lines used in this study were the mouse-derived striatal cells STHdh^Q7^ and STHdh^Q111^ (Coriell Cell Repositories). Cells were grown at 33°C in Dulbecco’s modified Eagle’s medium (DMEM, Genesee) supplemented with 10% fetal bovine serum (FBS), 100 U ml^-1^ penicillin/streptomycin and 100 ug ml^-1^ G418 (Gibco), as previously described (12).

### Mouse strains

For this study we used a full-length knock-in mouse model of HD known as zQ175 on the C57BL/6J background (Stock No. 027410). CK2α′ heterozygous mice (CK2α′^(+/−)^) on the 129/SvEv-C57BL/6J background (Taconic Biosciences TF3062) were originally obtained from Dr. Seldin (Boston University) (67). All mice were housed under standard SPF conditions. We also used 5-month WT (mixed background CBA x C57BL/6), R6/1, SNCA^KO^, and R6/1SNCA^KO^ obtained from Dr. Lucas. All animal care and sacrifice procedures were approved by the University of Minnesota Institutional Animal Care and Use Committee (IACUC) in compliance with the National Institutes of Health guidelines for the care and use of laboratory animals under the approved animal protocol 2007-38316A.

### siRNA transfection, RNA preparation and RT-qPCR

For CK2α’ knock-down, ST*Hdh* cells were transfected with FlexiTube siRNA (5 nmol) from Qiagen (Mm_Csnk2a2; SI00961051; SI00961058; SI00961065; SI00961072) using DharmaFECT1 per manufacturer’s guidelines. As a negative control, ON-TARGETplus control Non-targeting pool siRNA (Dharmacon) was used. Cells were collected 24 h after transfection. RNA was extracted from STHdh cells and mouse striatal tissues by using the RNeasy extraction kit (Qiagen) according to the manufacturer’s instructions. cDNA for all was prepared using the Superscript First Strand Synthesis System (Invitrogen). SYBR green based PCR was performed with SYBR mix (Roche). The qPCR amplification was performed using the LightCycler 480 System (Roche). Each sample was tested in triplicate and normalized to GAPDH levels.

### Immunoblot analysis

Sample preparation and immunoblotting condition were performed as previously described (12). Striatum protein extracts from one hemisphere of mice were prepared in cell lysis buffer (25 mM Tris pH 7.4, 150 mM NaCl, 1 mM EDTA, 1% Triton-X100 and 0.1% SDS). Primary antibodies were anti-CK2α’ (Novus NB100-379 and Proteintech 10606-1-AP), anti-Iba1 (FUJIFILM Wako 019-19741), α-syn (Biolegend 834303 clone 4D6), pS129-α-syn (Millipore MABN826, clone 81A and Abcam ab51253, EP1536Y), GAPDH (Santacruz sc-365062). Quantitative analyses were performed using ImageJ software and normalized to GAPDH controls.

### Immunohistochemistry

Sample preparation was performed as previously described (12). Fluorescent images were acquired on an epi-fluorescent microscope (Echo Revolve) or confocal microscope (Olympus FV1000). Primary antibodies used are as follows: α-syn (Biolegend 834303), pS129-α-syn (Millipore MABN826 and Cell signaling technology 23076S, D1R1R), CK2α’ (Proteintech 10606-1-AP), Ctip2 (Abcam ab18465), C3d (R&D Systems AF2655), GFAP (Invitrogen PA1-10019), S100b (Abcam ab41548), GS (BD Biosciences 610517 and Abcam 49873), HTT (Millipore, clone mEM48 Mab5374, and Abcam ab109115), Iba1 (FUJIFILM Wako 019-19741), NeuN (Millipore MAB377), IL-6 (Santa Cruz Bio sc-32296). For cell number (Ctip, GS, NeuN, Iba1, DAPI), the Cell counter plugin from ImageJ software was used and normalized to the image area (300μm^2^). EM48^+^ and α-syn puncta were counted using annotations in the Echo Revolve software and using the Puncta Analyzer plugin in ImageJ.

### Nuclear/Cytoplasm fractionation

Frozen striatum samples (∼20 mg) were fractionated using the Minute^TM^ Cytosolic and Nuclear Extraction Kit for Frozen/Fresh tissues (Invent Biotechnologies INC, Cat NT-032) as per Manufacturer’s instructions.

### Electrophysiological analyses

Acute dorsolateral striatum coronal slices (350 μm thick) were obtained from 12 months old mice using a vibratome, and processed as previously described (68). Researchers were blind to the mouse genotype. The brain was quickly removed after decapitation and placed in ice-cold artificial cerebrospinal fluid (ACSF) containing (in mM): NaCl 124, KCl 2.69, KH_2_PO_4_ 1.25, MgSO_4_ 2, NaHCO_3_ 26, CaCl_2_ 2, ascorbic acid 0.4, and glucose 10, and continuously bubbled with carbogen (95% O_2_ and 5% CO_2_) (pH 7.4). For excitatory postsynaptic currents (EPSCs) picrotoxin (50 µM) and CGP54626 (1 µM) were added. Whole-cell electrophysiological recordings were obtained using patch electrodes (3–10 MΩ) filled with an internal solution containing (in mM): KMeSO_4_ 135, KCl 10, HEPES-K 10, NaCl 5, ATP-Mg 2.5, GTP-Na 0.3 (pH 7.3). Membrane potentials were held at −70 mV. For EPSCs, theta capillaries filled with ACSF were used for bipolar stimulation. Input–output curves of EPSCs were made by increasing stimulus intensities from 0 to 100 μA. Paired-pulse facilitation was done by applying paired pulses (2 ms duration) with 25, 50, 75, 100, 200, 300, and 500 ms inter-pulse intervals. The paired-pulse ratio was calculated by dividing the amplitude of the second EPSC by the first (PPR=EPSC-2/EPSC-1). Synaptic fatigue was assessed by applying 30 consecutive stimuli in 15 ms intervals. For miniature EPSCs (mEPSCs) tetrodotoxin (TTX; 1 μM) was added to the solution.

### Behavioral assays

Sample sizes were calculated using GraphPad Prism 9.0 and GPower 3.1 to detect differences between WT versus zQ175 groups with a power of ≥ 0.8. Researchers at the Mouse Behavioral core at University of Minnesota were blinded to the genotypes of the mice during testing. See **Supplementary Methods** for a complete description of all behavioral tests conducted in the study. *Beam Walk*: 19-mm (medium-round) or 10-mm (small-round) diameter and 16-mm (medium-Square) or 10-mm (small-Square) width of 3 feet long wood beams (made in house) were used. Each mouse was placed on the beam at one end and allowed to walk to the goal box. Foot slips were recorded manually when the hind paws slipped off the beam. Testing included 3 training days and 1 test day with 4 consecutive trials each. *Rotarod*: Mice were tested over 3 consecutive days. Each daily session included 3 consecutive accelerating trials of 5 min on the rotarod apparatus (Ugo Basile) with the rotarod speed changing from 5 to 50 RPM over 300 s, with an inter-trial interval of at least 15 min.

### RNA-Seq Analyses

Gene expression analysis was carried out using the CHURP pipeline (>HTTps://doi.org/10.1145/3332186.3333156) using n=5 mice/genotype for WT, zQ175, and zQ175:CK2α’^(+/-)^ and n=3 mice for CK2α’^(+/-)^, with a female (F)/male (M) ratio: 4F/1M WT, 1F/2M CK2α’^(+/-)^, 2F/3M zQ175, 4F/1M zQ175:CK2α’^(+/-)^. Differential gene expression was determined with DESeq2 using default setting (PMID: 25516281). Genes with a q < 0.1 were considered significant. Outliers’ identification was performed using Cook’s distance (DESeq2). Driver factors of gene expression variance (genotype and/or sex) were evaluated using R package variance Partition. Pathway and clustering analysis were completed with IPA (Ingenuity Systems: RRID: SCR_008653) and gProfiler2 (PMID: 31066453). Data visualization was done using various R graphic packages, including ggplo2, ggraph, and DESeq2 visualization functions. The RNA-seq data set generated in this manuscript has been deposited at GEO (accession number GSE160586). Reviewer token “**gpqrigisbxgprqf”.**

### WGCNA Analysis

The count-based gene expressions were first transformed using a variance stabilizing method via DESeq2 vst function (69). The WGCNA R package (v1.69) was used to construct an unsigned gene co-expression network with a soft threshold power [beta] of 6. We used a non-parametric Kruskal-Wallis test (p value < 0.05) to identify modules that differed significantly among different genotypes. Data for the Greenyellow module was exported using a Cytoscape format for visualization. Network figures are limited to the top 15% of genes with the strongest network connections. The size of the circles is scaled by the absolute value of the mean log2 fold change between zQ175 and zQ175:CK2α’^(+/-)^ mice.

### Quantification and Statistical analyses

Data are expressed as Mean ± SEM, Mean ± SD, or percentage, analyzed for statistical significance, and displayed by Prism 8 software (GraphPad, San Diego, CA) or Excel software (Microsoft). Pearson correlation tests were applied to test the normal distribution of experimental data. Normal distributions were compared with Student t-test (two-tailed or one-tailed), Welch’s t-test or ANOVA with appropriate post-hoc tests (Sidak’s, Dunn’s, or Tukey’s) for multiple comparisons. The accepted level of significance was p ≤ 0.05. Statistical analyses for electrophysiological experiments were performed with SigmaPlot 13.0 software. No statistical methods were used to predetermine sample sizes, but sample sizes were chosen to be similar to those reported in previous publications (11).

## Acknowledgements

We are grateful to Drs. Sylvain Lesne and Michael Lee for sharing their expertise on alpha-synuclein and sharing reagents, Maha Syed and Joyce Meints for technical assistance, Jason Mitchell for assistance with confocal microscopy and Erin Greguske for proofreading.

## Authors’ contributions

R.G.P obtained funding for the study and designed the experiments. D.Y, N.Z, F.C, J.Y, T.B, W.T, K.J, T.S.M, K.G, S.L, A.W, D.C, R.M performed the experiments. C.T.Z prepared SNCA^KO^ tissues. Y.Z conducted RNA-seq analyses. D.Y, N.Z, R.M, S.L, K.G, C.N, W.T and R.G.P prepared and analyzed the data. G.O supervised the MR data acquisition and analysis. M.B supervised mouse behavioral data analysis. A.A supervised electrophysiological recordings. M.C supervised microglia analyses. J.J.L supervised SNCA^KO^ tissue preparation. R.G.P wrote the first draft of the manuscript and all authors edited subsequent versions and approved the final version of the manuscript.

## Data availability

RNA-seq data set generated in this manuscript is accessible at GEO (accession number GSE160586). All other data generated or analyzed during this study are included in this published article (and its supplementary information files).

