## Supplementary Information for "CK2 alpha prime and alpha-synuclein pathogenic functional interaction mediates synaptic dysregulation in Huntington’s disease"

### Supplementary Methods

### Cell lines

Mouse-derived striatal cells STHdh$^{Q7}$ and STHdh$^{Q111}$ (Coriell Cell Repositories) were grown at 33ºC in Dulbecco's modified Eagle's medium (DMEM) supplemented with 10% fetal bovine serum (FBS), 100 U ml$^{-1}$ penicillin/streptomycin and 100 ug ml$^{-1}$ G418 (Gibco), as previously described (11).

### Mouse strains

All reported data abides to the ARRIVE guidelines for reporting animal research. For this study we used a full-length knock-in mouse model of HD known as zQ175 on the C57BL/6J background (B6J.zQ175 KI mice (Stock No. 027410)), which harbors a chimeric human/mouse exon 1 carrying an expansion of ~188 CAG repeats and the human poly-proline region (27). WT (C57BL/6) animals were used as controls. CK2α′ heterozygous mice (CK2α′$^{(+/-)}$) on the 129/SvEv-C57BL/6J background (Taconic Biosciences TF3062) were originally obtained from Dr. Seldin (Boston University) (71). CK2α′$^{(+/-)}$ were crossbred with WT and zQ175 for more than 10 generations prior to selecting the founder animals of the study. All mice used have the C57BL/6J background. All mice were housed under standard SPF conditions in a conventional 12:12 *light*/*dark* cycle. All experiments were conducted during the light cycle. Mice were bred as previously described [11] to generate the following genotypes: WT (CK2α′$^{(+/+)}$ HTT$^{(0/0)}$), CK2α′$^{(+/-)}$ (CK2α′$^{(+/-)}$ HTT$^{(0/0)}$), zQ175 (CK2α′$^{(+/+)}$ HTT$^{(Tg/0)}$), zQ175;CK2α′$^{(+/-)}$ (CK2α′$^{(+/-)}$ HTT$^{(Tg/0)}$). Animals were analyzed at 3, 6, 12, and 22 months of age. We also used samples from 5 month WT (mixed background CBA x C57BL/6), R6/1, SNCA$^{KO}$, and R6/1SNCA$^{KO}$ obtained from Dr. Lucas. Sample size was set to a minimum of three animals per genotype for every analysis. When possible, at least two females and two males were used for each genotype and age. Littermates of the same sex were randomly assigned to experimental groups. All animal care and sacrifice procedures were approved by the University of Minnesota Institutional Animal Care and Use Committee (IACUC) in compliance with the National Institutes of Health guidelines for the care and use of laboratory animals under the approved animal protocol 2007-38316A.

**SNCA siRNA or overexpressing plasmid transfection**

For SNCA knock-down, ST*Hdh* cells were transfected with FlexiTube siRNA (1 nmol) from Qiagen (Ms_SNCA; SI04945570;SI04945563;SI04945556;SI00183155) using DharmaFECT1 per manufacturer's guidelines. As a negative control, we used scramble siRNA-A (Santa Cruz sc-37007). For SNCA overexpression EGFP WT-alpha synuclein plasmid (#40822) from Addgene was transfected using 3 ug. As a control, cells were transfected with 3ug of a pEGFP-C1 plasmid (Clontech). Cells were collected 24 h after transfection. RNA was extracted from STHdh cells and mouse striatal tissues by using the RNeasy extraction kit (Qiagen) according to the manufacturer's instructions.

**RNA preparation and RT-qPCR**

RNA was extracted from STHdh cells and mouse striatal tissues by using the RNeasy extraction kit (Qiagen) according to the manufacturer's instructions. cDNA for all samples was prepared from 1 µg RNA using the Superscript First Strand Synthesis System (Invitrogen) according to the manufacturer's instructions. SYBR green based PCR was performed with SYBR mix (Roche). The primer sequences for target genes were as follows: CK2α' forward: 5'- CGACTGATTGATTGGGGTCT-3' reverse: 5'-AGAATGGCTCCTTTCGGAAT-3', IL-6 forward: 5'- AGTTGCCTTCTTGGGACT-3' reverse: 5'-TCCACGATTTCCCAGAGAAC-3', PSD-95 (Dlg4) forward: 5'-CCGACAAGTTTGGATCCTGT-3', reverse: 5'-ACGGATGAAGATGGCGATAG, Drd1 forward: 5'-AAGATGCCGAGGATGACAAC-3', reverse:5'-CCCTCTCCAAAGCTGAGATG-3', Drd2 forward: 5'-TATGCCCTGGGTCGTCTATC-3', reverse: 5'-AGGACAGGACCCAGACAATG-3', Darpp32(PPP1R1B) forward: 5'-CCACCCAAAGTCGAAGAGAC-3', reverse: 5'-GCTAATGGTCTGCAGGTGCT-3', GAPDH (used as an internal control gene) forward: 5'-AACTTTGGCATTGTGGAAGG-3' reverse: 5'-ACACATTGGGGGTAGGAACA-3'. The qPCR amplification was performed using the LightCycler 480 System (Roche). Each sample was tested in triplicate and normalized to GAPDH levels. For analysis, the $2^{\Delta\Delta Ct}$ method was used to calculate the relative fold gene expression of samples.

**Immunoblot analysis**

Sample preparation and immunoblotting were performed as previously described (11). Protein extracts were prepared in cell lysis buffer (25 mM Tris pH 7.4, 150 mM NaCl, 1 mM EDTA, 1% Triton-X100 and 0.1% SDS). Primary antibodies were anti-CK2α' (Novus NB100-379 and Proteintech 10606-1-AP), anti-Iba1 (FUJIFILM Wako 019-19741), α-syn (Biolegend 834303 clone 4D6), pS129-α-syn (Millipore MABN826, clone 81A and EP1536Y Abcam ab51253), GAPDH (Santacruz sc-365062).

#### Immunohistochemistry

Mice were anesthetized with Avertin (250 mg/kg Tribromoethanol) and perfused intracardially with 25mM Tris-base, 135mM Nacl, 3mM KCl, pH 7.6 supplemented with 7.5 mM heparin, as previously described (11). Brains were dissected, fixed with 4% PFA in TBS at 4°C for 4-5 days, cryoprotected with 30% sucrose in TBS for 4-5 days and embedded in a 2:1 mixture of 30% sucrose in TBS:OCT (Tissue-Tek). Immunostaining was performed as previously described (11). Sample preparation was performed as previously described (11). Fluorescent images were acquired on an epi-fluorescent microscope (Echo Revolve) or confocal microscope (Olympus FV1000). Primary antibodies used and dilutions are as follows: α-syn (Mouse, Biolegend 834303, 1:1000), pS129-α-syn (Mouse, Millipore MABN826, 1:500 and Rabbit, D1R1R Cell signaling technology 23076S, 1:200), CK2α' (Rabbit, Proteintech 10606-1-AP, 1:500), Ctip2 (Rat, Abcam ab18465, 1:500), complement component C3d (Goat, R&D Systems AF2655, 1:200), Glial fibrillary acidic protein GFAP (Rabbit, Invitrogen PA1-10019, 1:500), S100 Calcium Binding Protein B S100b (Rabbit, Abcam ab41548, 1:500), glutamine synthetase GS (Mouse, BD Biosciences 610517, 1:1000 and Rabbit, Abcam 49873, 1:500), HTT (Mouse, Millipore, clone mEM48 Mab5374, 1:100, and Rabbit, Abcam ab109115, 1:500), Iba1 (Rabbit, FUJIFILM Wako 019-19741, 1:200), NeuN (Mouse, Millipore MAB377, 1:1000), IL-6 (Mouse, Santa Cruz Bio sc-32296, 1:100). For cell number (Ctip, GS, NeuN, Iba1, DAPI) the Cell counter plugin from ImageJ software was used and normalized to the image area ($300 \mu m^2$). $EM48^+$ aggregates were counted using annotations in the Echo Revolve software and validated using the Puncta Analyzer plugin in ImageJ. Number of pS129-α-syn/EM48 puncta was calculated using the Puncta Analyzer plugin in ImageJ.

#### Cytokine and chemokine array

The Proteome Profiler Mouse Cytokine Array Panel (ARY006, R&D Systems) was used to detect the levels of cytokine/chemokine of the mouse brain as per manufacturer's instructions. Frozen striatum from 12 - 14 months was homogenized in PBS containing Halt protease inhibitor cocktail and phosphatase inhibitors (Fisher Scientific) and 1% triton X-100 (Sigma). Samples were stored at -80°C for 15 min, thawed and centrifuged at 10,000 × g for 5 min to remove cell debris. A total of n=6 mice/genotype with a female(F)/male(M) ratio (3F/3M WT, 5F/1M zQ175, and 4F/2M zQ175:CK2α'$^{(+/-)}$) were analyzed. Samples were grouped in three different pools for each genotype: pools 1 and 2 contained three different mice per genotype, pool 3 contained a randomized selection of 3 mice/genotype out the n=6 mice cohort. Similar numbers of male and female mice were evaluated. Data was analyzed using ImageJ software and presented as an average signal of three independent pool assays, calculating pairs of duplicate spots corresponding to 40 different cytokines or chemokines.

#### Cresyl Violet staining

Coronal brain slices were mounted on Fisherbrand Superfrost Plus Microscope Slide and dried at 37 ºC overnight. Dried sections were incubated with 0.2% Cresyl violet acetate (Sigma #C5042) pH 3.7 for 8 minutes. Samples were then dehydrated in 80%,

95%, and 100% Ethanol solution for two minutes each followed by Xylene incubation. Slides were dried and mounted with permount (#SP15-500) and imaged on a Leica DM2500. Neurons were identified as round light blue cells and manually counted.

**Electrophysiological analyses**

Acute dorsolateral striatum coronal slices (350 µm thick) were obtained from approximately 12 months old mice using a vibratome, and processed as previously described (72). Researchers were blind to the mouse genotype. The brain was quickly removed after decapitation and placed in ice-cold artificial cerebrospinal fluid (ACSF). Slices were incubated for at least 1h before use in a chamber at room temperature (21–24 ºC) in ACSF containing (in mM): NaCl 124, KCl 2.69, $KH_2PO_4$ 1.25, $MgSO_4$ 2, $NaHCO_3$ 26, $CaCl_2$ 2, ascorbic acid 0.4, and glucose 10, and continuously bubbled with carbogen (95% $O_2$ and 5% $CO_2$) (pH 7.4). Slices were then transferred to an immersion recording chamber and superfused at 2 mL/min with gassed ACSF at 30-32 ºC and visualized under an Olympus BX50WI microscope (Olympus Optical; Japan). To study excitatory postsynaptic currents (EPSCs) picrotoxin (50 µM) and CGP54626 (1 µM) were added to the solution to block the $GABA_A$ and $GABA_B$ receptors, respectively. Whole-cell electrophysiological recordings were obtained from medium spiny neurons (MSNs) using patch electrodes (3–10 MΩ) filled with an internal solution containing (in mM): $KMeSO_4$ 135, KCl 10, HEPES-K 10, NaCl 5, ATP-Mg 2.5, GTP-Na 0.3 (pH 7.3). Recordings were obtained with a PC-ONE amplifier (Dagan Instruments; Minneapolis, MN, USA). Membrane potentials were held at −70 mV. Signals were filtered at 1 kHz, acquired at a 10 kHz sampling rate, and fed to a Digidata 1440A digitizer (Molecular Devices; San Jose, CA, USA). pCLAMP 10.4 (Axon Instruments, Molecular Devices; San Jose, CA, USA) was used for stimulus generation, data display, data acquisition, and data storage. To record evoked EPSCs, theta capillaries filled with ACSF were used for bipolar stimulation and placed in the vicinity of the cell patched in the dorsolateral striatum. Input–output curves of EPSCs were made by increasing stimulus intensities from 0 to 100 µA (WT, n = 8; zQ175, n = 9; zQ175:CK2α'$^{(+/-)}$ n = 13). Paired-pulse facilitation was done by applying paired pulses (2 ms duration) with 25, 50, 75, 100, 200, 300, and 500 ms inter-pulse intervals. The paired-pulse ratio was calculated by dividing the amplitude of the second EPSC by the first (PPR=EPSC-2/EPSC-1). Synaptic fatigue was assessed by applying 30 consecutive stimuli in 15 ms intervals. For miniature EPSCs (mEPSCs) tetrodotoxin (TTX; 1 µM) was added to the solution. Amplitud and frequency was analyzed from WT, n = 10; zQ175, n = 9; zQ175:CK2α'$^{(+/-)}$ n = 12). Short term potentiation was analyzed from WT, n = 8; zQ175, n = 9; zQ175:CK2α'$^{(+/-)}$ n = 11. Short term depression was analysed from WT, n = 7; zQ175, n = 9; zQ175:CK2α'$^{(+/-)}$ n = 12.

**RNA-Seq Analyses**

Gene expression analysis was carried out using the CHURP pipeline (HTTps://doi.org/10.1145/3332186.3333156) using n=5 mice/genotype for WT, zQ175, and zQ175:CK2α'$^{(+/-)}$ and n=3 mice for CK2α'$^{(+/-)}$, with a female (F)/male (M) ratio: 4F/1M WT, 1F/2M CK2α'$^{(+/-)}$, 2F/3M zQ175, 4F/1M zQ175:CK2α'$^{(+/-)}$. Read quality was assessed using FastQC (HTTp://www.bioinformatics.bbsrc.ac.uk/projects/fastqc). Raw reads were then trimmed to remove low quality 3' ends and adaptor contamination using Trimmomatic with the following parameter: ILLUMINACLIP:all_illumina_adapters.fa:4:15:7:2:true LEADING:3 TRAILING:3 SLIDINGWINDOW:4:15 MINLEN:18 (74). Processed reads were aligned to mouse reference genome GRCm38 via HiSat2 (PMID: 31375807) using default parameters. Post-alignment cleaning removed duplicated mapping, unmapped reads, and reads with MAPQ<60. Gene-level expressions were quantified using Subread (PMID: 24227677), and differential gene expression was determined with DESeq2 using default setting (PMID: 25516281). Genes with a q < 0.1 were considered significant. Outliers' identification was performed using Cook's distance (DESeq2), which is a measure of how much a single sample is influencing the fitted coefficients for a gene. Driver factors of gene expression variance (genotype and/or sex) were evaluated using R package variance Partition. Pathway and clustering analysis were completed with Ingenuity Pathway Analysis (Ingenuity Systems: RRID: SCR_008653) and gProfiler2 (PMID: 31066453). Data visualization was done using various R graphic packages, including ggplo2, ggraph, and DESeq2 visualization functions. The RNA-seq data set generated in this manuscript has been deposited at GEO (accession number GSE160586). Reviewer token "**gpqrigisbxgprqf".**

**WGCNA Analysis**

Genes with less than 10 counts in more than 90% of samples were removed from WGCNA analysis (75). The count-based gene expressions were first transformed using a variance stabilizing method via DESeq2 vst function. The WGCNA R package (v1.69) was used to construct an unsigned gene co-expression network with a soft threshold power [beta] of 6. Nineteen tightly connected modules were detected, while genes that were not connected to the 19 modules were collected in the $20^{th}$ "grey" module. WGCNA defines the expression of a given module by averaging module gene expressions, which is also called module eigengene expression. We used a non-parametric Kruskal-Wallis test (p value < 0.05), to identify modules that differed significantly among mouse samples with different genotypes. For example, the "Greenyellow" module differs significantly between zQ175 mice and zQ175:CK2α'$^{(+/-)}$ mice. Data for the Greenyellow module were exported using a Cytoscape format for visualization. Network figures are limited to the top 10% of genes with the strongest network connections (the topological overlap threshold was raised until only 10% of genes remained). The network modules are color coded by the differential expression between zQ175 mice and zQ175:CK2α'$^{(+/-)}$ mice: blue, downregulated in zQ175 compared to ZQ175:CK2α'$^{(+/-)}$ mice; red, upregulated in zQ175 compared to ZQ175:CK2α'$^{(+/-)}$. The size of the circles is scaled by the absolute value of the mean log2 fold change between zQ175 and ZQ175:CK2α'$^{(+/-)}$ mice.

**Measurement of striatal volume**

For volumetric analysis of striatum, coronal RARE images were used. The images had an in-plane resolution of 0.125 mm x 0.125 mm and 1 mm slice thickness. All volumetric quantifications were performed using ImageJ. The perimeter of the striatum was traced using a polygon selection tool and the volume measurement was performed by running

Measure Stack plugin on ImageJ. All quantitative analyses were conducted blindly with respect to mouse genotype.

**Behavioral assays**

Sample sizes were calculated using GraphPad Prism 9.0 and GPower 3.1. and determined a minimum sample size of 6 mice per group to detect differences between WT versus zQ175 groups with a power of ≥ 0.8.

*Barnes Maze*: The Barnes maze is a dry-land maze test for spatial learning and memory. It consists of a circular platform with 20 holes along the perimeter (San Diego Instruments). Spatial cues were placed on all four walls of the behavioral testing room, lit to 300 lx during testing. Mice were acclimatized to the testing room for 30 min at the beginning of each training session. Training days consisted of 4 trials per day for 4 days. Training trials ended when the subject climbed into the escape box within the goal quadrant located under 1 of the holes or when the maximum trial duration of 180 s was reached. Subjects were run in small groups of six mice or less, so that no more than 20 min passed between trials for a given animal during training. On the day following the last training trial, memory was assessed in single 90 s probe trial tests, where the target escape box in the goal quadrant was replaced with a false box cover identical to the other 19 holes, and the exploration pattern of each subject was examined.

*Barnes Maze Reversal*: The Barnes Maze Reversal was conducted in the same way as Barnes Maze as described above besides two differences: Test consisted of 2 training days as opposed to 4 in the Barnes Maze, and the escape hole was located on the opposite side of the maze.

*Cylinder Test*: Mice were placed in a transparent glass cylinder (40 cm length and 20 cm in diameter, made in house). The number of forelimb contacts while rearing against the wall of the cylinder was scored manually for 5 mins to examine asymmetric motor impairments.

*Delay Fear Conditioning*: The test was conducted in a chamber (Interior: 23.5"x 28"x 12.5", Exterior: 25"x 29.5"x14") equipped with a speaker and a stainless-steel foot shock grid floor (Med Associates Inc.). On Day 1, mice were trained on a 9-min training session in a testing chamber. Then, animals received 5 paired light (15-20 lx) and tone cues (8000Hz, 80 dB; Rise/fall 10ms) immediately followed by a foot shock (0.7 mA), with 60 second-intertrial intervals. Mice were then removed from the chamber and returned to its home cage. 24 h after training (Day 2), mice were re-exposed to the test chamber (no cue presentation) for 3 mins and context-dependent freezing was recorded for baseline freezing measurement. Then, mice spent 6 mins in a test chamber which presents paired light and tone cues in the last 2 mins of testing to induce cued freezing. Data collection and analysis were semi-automated via video-monitoring fear-conditioning apparatus (Med Associates, Inc.).

***Open Field Test***: The pattern of exploration is used as a measure of anxiety-like behavior. Activity chambers (20”x20”x10”, made in house) were equipped with light intensity of 250 lx. Mice were placed in the center of the chamber and their behavior was recorded using ANY-maze software for 30 mins. The test arena was divided into a center and periphery zone. The periphery was a ~3.5-inch-wide zone adjacent to the walls and was used to determine thigmotactic behavior. Analyses were performed on the following five measures: total locomotion (distance traveled), locomotion in the inner or outer zone, and velocity in the light phase of the cycle.

*Y-Maze*: This test evaluated short-term spatial and working memory on a Y-maze apparatus (made in house) with three arms (15.75” × 3.5” × 7.25”) at a 120 ° angle. Animals were first moved to a testing room and habituated for 15-30 mins while setting up the apparatus. To measure spatial short-term memory, one of the three arms of the maze (“Novel Arm”) was blocked and the mouse was placed in the start arm, allowed to explore the other two arms for 10 mins while the time spent in each zone was recorded. Then, the mouse was transferred to its cage for 60-min of inter-trial interval before the test trial. Mice were then returned to the initial arm and allowed to wander around all three arms of the maze for 5 mins. Time spent in each arm and the number of entries into each arm were recorded. Spontaneous alternation task was used to assess spatial working memory and was performed at least 20 hours after the spatial memory task. In this task, the innate response of a mouse to a new environment was evaluated. Alternation was defined as entry into all three arms, e.g., ABC, BCA, or CAB, but not CAC. The mouse was habituated in the testing room for 15-20 mins while setting up the apparatus. The animal was then placed in one arm of the maze and was allowed to move freely all three arms for 5 minutes. Latency to exit the start arm, the number and pattern of arm choices were recorded.

**Magnetic resonance imaging and spectroscopy**

A quadrature surface radio frequency (RF) coil with two geometrically decoupled single turn coils (14 mm diameter) was used as the MR transceiver. Following positioning of the mouse in the magnet, coronal and sagittal multislice images were obtained using a rapid acquisition with relaxation enhancement (RARE) sequence [repetition time (TR)= 4 s, echo train length= 8, echo time (TE)= 60 ms, slice thickness= 1 mm, 7 slices]. The volume of interest (VOI) studied was centered on the striatum (8.2 μl, 1.7 x 2.0 x 2.4 $mm^3$). All first- and second-order shims were adjusted using FASTMAP with echo-planar readout. Localized $^1H$ MR spectra were acquired with a short-echo localization by adiabatic selective refocusing (LASER) sequence [TE= 15 ms, TR= 5 s, 256 transients] combined with VAPOR (variable power RF pulses with optimized relaxation delays) water suppression. Spectra were saved as single scans. Unsuppressed water spectra were acquired from the same VOI for metabolite quantification. *Metabolite quantification*; Single shots were eddy current, frequency, and phase corrected using MRspa software (HTTp://www.cmrr.umn.edu/downloads/mrspa/) before averaging. The contributions of individual metabolites to the averaged spectra were quantified using LCModel as described previously[83]. The following metabolites were included in the basis set: alanine (Ala), ascorbate/vitamin C (Asc), aspartate (Asp), glycerophosphocholine (GPC), phosphocholine (PCho), creatine (Cr), phosphocreatine (PCr), gamma-aminobutyric acid (GABA), glucose (Glc), glutamine (Gln), glutamate (Glu), glutathione (GSH), glycine (Gly), *myo*-inositol (Ins), lactate (Lac), *N*-acetylaspartate (NAA), *N*-acetylaspartylglutamate

(NAAG), phosphoethanolamine (PE), *scyllo*-inositol (Scyllo), taurine (Tau), and macromolecules (MM). The MM spectra were experimentally obtained from a VOI that covered the striatum using an inversion recovery technique [VOI = 4.7 x 2.1 x 2.7 $mm^3$, TE = 15 ms, TR = 2 s, inversion time (TIR) = 675 ms, 400 transients, N = 2]. The model metabolite spectra were generated using density matrix simulations with the MATLAB software (MathWorks) based on previously reported chemical shifts and coupling constants[84]. Concentrations with mean Cramér-Rao lower bounds ≤20% in any of the 3 groups were reported. If the correlation between two metabolites was consistently high (correlation coefficient $r < -0.7$), their sum was reported rather than the individual values[83]. Strong negative correlation was found in two cases, so that Cr and PCr (denoted tCr for total creatine) and GPC and PCho (denoted tCho for total choline) were reported.

**Quantification and Statistical analyses**

Data are expressed as Mean ± SEM, Mean ± SD, or percentage, analyzed for statistical significance, and displayed by Prism 8 software (GraphPad, San Diego, CA) or Excel software (Microsoft). Pearson correlation tests were applied to test the normal distribution of experimental data. Normal distributions were compared with t-test (two-tailed or one-tailed) or ANOVA with appropriate post-hoc tests (Sidak's, Dunn's, or Tukey's) for multiple comparisons. Non-normal distributions were compared with the non-parametric Kruskal-Wallis test with an appropriate post-hoc test, as indicated. The accepted level of significance was $p \leq 0.05$, therefore samples indicated as n.s (no significant) did not reach $p \leq 0.05$. Statistical analyses for electrophysiological experiments were performed with SigmaPlot 13.0 software. No statistical methods were used to predetermine sample sizes, but sample sizes were chosen to be similar to those reported in previous publications (11). In the figures, we show the mean as an estimator of central tendency including when we have used a non-parametric test, to maintain consistency with other figures in the paper and because it is more intuitive to compare the mean values.

**Supplementary Figure Legends**

**Figure S1**. **Depletion of MSN marker expression does not reflect neuronal loss in zQ175 mice**. ***a***, Representative images of EM48 immunostaining in the dorsal striatum and cortex of zQ175 at 3, 6, 12 and 22 months (*n*=6 mice/genotype). ***b***, Quantification of the number of EM48 aggregates in the cortex of zQ175 and its comparison with striatum. ***c***, Representative images of NeuN immunostaining in the dorsal striatum of WT and zQ175 at 3, 6, 12 and 22 months (*n*=3 mice/genotype). **d*, e***, Representative images of Ctip2 immunostaining in the dorsal striatum of 12 months old WT and zQ175 and quantitative analysis of Ctip2+ cells (*n* = 4 WT; 5 zQ175). Scale bar, 50 µm. DAPI is used for nuclear staining. ***f***, Cresyl violet staining in the dorsal striatum of 22 months old WT and zQ175 mice. Magnified image represents neurons (white arrow) and glial (black arrow) cells. ***g,*** Quantification of neurons from cresyl violet analyses at 12 and ***h***, 22 months old WT and zQ175 mice. At least 3 mice/genotype and 3 images per mouse were analyzed. ***i***, Dotted line on magnetic resonance images displays the manually traced striatum region of representative mice of each genotype at 22 months of age. ***j***, Striatum volume analyzed in 22-month-old mice from images in G (n=4 mice/genotype). Error bars denote mean ± SD. Student's t-test, p-values <0.05 are indicated. n.s = not significant.

**Figure S2**. Deficiency in CK2α' expression does not affect MSNs but does influence synaptic proteins. ***a, b***, Representative images show the labeling of Ctip2 (***a***) and quantitative analysis (**b**) from striatum of zQ175 and zQ175:CK2α'$^{(+/-)}$ mice (*n*=3 mice/genotype). Scale bar, 50 µm. ***c***, mRNA levels assessed by RT-qPCR of Drd1 and Drd2 (striatal MSN markers), Darpp32 and PSD95 in the striatum of 12 months old mice (*n*=4 mice/genotype). Error bars denote mean ± SD, values were analyzed by Student's t-test. *p*-values <0.05 for differences between groups are indicated in each graph. n.s = not significant.

**Figure S3**. Genetic deletion of CK2α' does not ameliorate anxiety-like behaviors or spatial working memory of zQ175 HD at 12 months of age. **a-*e***, Gross motor performance, exploratory behavior, and anxiety-like behavior were assessed during 30 mins in an open field test. **a**, distance traveled, ***b***, time spent at the center of the field, ***c***, average locomotion velocity and ***d***, time in the outer/inner zone of the field between zQ175 and zQ175:CK2α' $^{(+/-)}$. ***e***, Representative tracing images show the total distance traveled by the subject. ***f***, Parameters of the spontaneous motor activity (number of rears) evaluated by a cylinder test. Error bars denote mean ± SEM, values were analyzed by two-way ANOVA with *Sidak's* post-hoc test. n.s = not significant.

**Figure S4**. CK2α' does not alter cognitive behavior in symptomatic HD mice. ***a***, ***b***, Freezing time in the cued (***a***) and contextual fear conditioning test (***b***). ***c-e***, Performance on the Barnes maze (BM) task measured by latency to reach the escape hole (***c***), the mean distance from the target location (***d***), and total distance until escape in training sessions (***e***). ***f-h,*** Results of BM reversal test are similar to those in BM test, measured by latency to first entry (***f***), total distance traveled (***g***) and the mean distance from the target location (***h***). ***i***, working memory, percent of spontaneous alternation measured by Y maze. Tests were conducted in 12 month old zQ175 and zQ175:CK2α'$^{(+/-)}$.Error bars represent mean ± SEM. Statistical analyses were conducted using ANOVA with *Sidak's* post-hoc test (*n* = 6 mice/genotype). n.s = not significant.

**Figure S5**. Expressions of all identified co-expression gene modules from WGCNA studies for each mouse sample. A total of 20 different modules were identified when comparing all the genotypes.

**Figure S6**. **RNA-Seq comparison between samples and with Langfelder et al., 2016 (35).** ***a***, Kruskal-Wallis test of module expressions between zQ175 mice and WT mice. ***b*** IPA canonical pathway analysis of module genes for module "Red". **c**, Cook's distance (DESeq2), which is a measure of how much a single sample is influencing the fitted coefficients for a gene, for all tested samples (HET= CK2α'$^{(+/-)}$, KI=zQ175, KIHET= zQ175:CK2α'$^{(+/-)}$). ***d,*** Network visualization of top 15% connected genes for module genes of module "Greenyellow". The size of the circles was scaled by the absolute value of the mean log2-fold change between zQ175 and zQ175:CK2α'$^{(+/-)}$ mice. **e**, MA-plot of differential gene expression between HD (zQ175) and WT mice. ***f,*** Venn Diagram of differentially expressed genes between HD (zQ175) and WT mice in comparison to Langfelder 2016 data. ***g,*** MA-plot of differential gene expression between zQ175:CK2α'$^{(+/-)}$ and WT mice. ***h,*** Venn Diagram of differentially expressed genes between zQ175, WT, zQ175:CK2α'$^{(+/-)}$ and CK2α'$^{(+/-)}$ mice. ***i***, Driver factors of gene expression variance (genotype and/or gender) for the DGEs identified between zQ175 and zQ175:CK2α'$^{(+/-)}$ were evaluated using R package variance Partition.

**Figure S7**. Alteration in CK2α' expression changes cytokine profiles but not microglia. **a**, RT-qPCR analysis for CK2α' and IL-6 in ST*Hdh*$^{Q7/Q7}$ (control) and ST*Hdh*$^{Q111/Q111}$ (HD) cell (n = 5 independent experiments). Data were normalized to GAPDH and control cells. **b**, siRNA knockdown of CK2α' for 24 h in ST*Hdh*$^{Q111/Q111}$ cells and RT-qPCR. Data were normalized with GAPDH and relativized to non-targeting control siRNA-treated cells (scramble), (*n* = 6 independent experiments). ***c, d,*** Representative images of mouse cytokine array panels from striatum extracts of WT, zQ175, and zQ175:CK2α'$^{(+/-)}$ at 12-14 months of age (*n* = 6 mice/genotype). **e**, **f**, Iba1 immunoblotting in the striatum of 12 months old WT, zQ175 and zQ175:CK2α', quantification was measured by image analyses using Image J software, GAPDH is used as a loading marker. **g-i**, Representative images show the labeling of Iba1 in the dorsal striatum of WT, zQ175 and zQ175:CK2α'$^{(+/-)}$ at 3, 6 and 12 months old (**g**). Images were analyzed using the Image J software. Scale bar, 100 μm. Iba1$^+$ cell body area in 12 months old mice (n=4 mice/genotype) (**h**) and percent of Iba1$^+$ cells in 300mm$^2$ (**i**). Error bars denote mean ± SEM, values were analyzed by Student's t test in **d**, and one-way ANOVA and Tukey post-hoc test in **a**, **b**, **f**, **h**, **i**. **p* represent p-values comparing zQ175 and WT, #*p* are p-values comparing zQ175 and zQ175:CK2α'$^{(+/-)}$.

**Figure S8**. Decreased CK2α' ameliorated astrogliosis in zQ175 mice. **a**, **b**, Representative images show the labeling (**a**) and quantification (**b**) of Glutamine synthetase (GS, an astrocytic marker) and CK2α' of striatum sections from 12-month-old mice in WT, zQ175 and zQ175:CK2α'$^{(+/-)}$ (*n* = 5 mice/genotype). DAPI is used for nuclear staining. Scale bars, 50 μm. **c**, Coronal images of brain scans in 9.4T magnet showing the striatum voxel (green box) for MRS acquisition from each genotype at 22 months. **d**, Localized proton magnetic resonance spectra [LASER sequence, TE= 15 ms, TR= 5 s, 256 transients, 9.4T] obtained at 22 months of age from the striatum of WT, zQ175, and zQ175:CK2α'$^{(+/-)}$ mice (*n* = 4 WT; 4 HD; 6 HD: CK2α' $^{(+/-)}$). Differences in *myo*-inositol (Ins) between WT and zQ175 mice and between zQ175 and zQ175:CK2α'$^{(+/-)}$ mice are shown with arrows. **e**, **f**, Mean concentrations of *myo*-inositol (Ins) (**e**) and other reliably quantified metabolites (**f**) in the striatum of WT (black), zQ175 (red), and zQ175:CK2α'$^{(+/-)}$ (gray) mice. Asc: Ascorbate/vitamin C; tCr: total creatine + phosphocreatine; GABA: gamma-aminobutyric acid; Glc: glucose; Gln: glutamine; Glu: glutamate; tCho: total phosphocholine + glycerophosphocholine; GSH: glutathione; Ins: *myo*-inositol; Lac: lactate; NAA: *N*-acetylaspartate; NAAG: *N*-acetylaspartylglutamate; PE: phosphoethanolamine; Tau: taurine. Error bars denote mean ± SEM, values were analyzed by one-way ANOVA with Tukey's post-hoc test. *p*-values < 0.05 are indicated.

**Figure S9.** SNCA regulates the expression of synaptic genes identified in IPA analysis in R6/1 mice. **a,** IB for α-syn (4D6 antibody) and mtHTT (EM48 antibody) in the striatum of 5 months old WT, SNCA$^{KO}$, R6/1 and R6/1:SNCA$^{KO}$ (n=3 mice/genotype). ***b***, RT-qPCR analyses for SNCA and genes identified in the IPA analyses to be connected to SNCA; (Ttr, Grm2, Slc17a7, Slc30a3, Cckbr, Nrp2, Tbr1 and Nr4a2) (n=4-5 mice/genotype). Error bars represent mean ± SEM. Statistical analyses were conducted by Student's t-test. p-values <0.05 are indicated. We also indicated p-values <0.09 for those genes that showed a trend toward decreased expression.

**Figure S10.** CK2a' haploinsufficiency decreased the levels of pS129-α-syn. ***A***, Representative pS129-α-syn IF images (D1R1R antibody) in the dorsal striatum of 12 months old WT, zQ175 and zQ175:CK2α'$^{(+/-)}$, Scale bar, 20 µm. ***B***, pS129-α-syn and CK2α' immunoblotting of striatum samples from 12-month-old zQ175 and zQ175:CK2α'$^{(+/-)}$ mice (n=3 mice/group). ***C***, Levels of pS129-α-syn and CK2α' were calculated using Image J from immunoblotting images in ***A*** and showed a parallel decrease of pS129-α-syn and CK2α' levels in zQ175:CK2α'$^{(+/-)}$ compared to zQ175 mice. Error bars represent mean ± SEM. Statistical analyses were conducted by Student's t-test. p-values <0.05 are indicated (*n*=3 mice/genotype).

**Dataset S1.** WGCNA modules genes assignment.

**Dataset S2.** WGCNA module names and number of genes per module.

**Dataset S3.** Differential Gene Expression analyses across the four different genotypes WT, zQ175, CK2α'$^{(+/-)}$ and zQ175:CK2α'$^{(+/-)}$.

**Dataset S4.** Expression analyses for astrocyte markers of A1, A2 and pan-reactive (from Liddelow et al., 2017) in WT, zQ175, CK2α'$^{(+/-)}$ and zQ175:CK2α'$^{(+/-)}$.

**Dataset S5.** Expression analyses for microglia markers of A1 inducing and reactive microglia (from Liddelow et al., 2017 (40)) in WT, zQ175, CK2α'$^{(+/-)}$ and zQ175:CK2α'$^{(+/-)}$.

**Dataset S6.** Marker genes and their mean log2 fold change between zQ175 and zQ175:CK2α'$^{(+/-)}$ mice compared to WT.

**Supplementary Movie 1**. Identification of pS129-α-syn nuclear localization (red) in the striatum of 12-month-old zQ175 mice. DAPI (bue) stains nuclei.
