## Supplementary figures and images for "CK2 alpha prime and alpha-synuclein pathogenic functional interaction mediates synaptic dysregulation in Huntington’s disease"

### Supplementary Figures

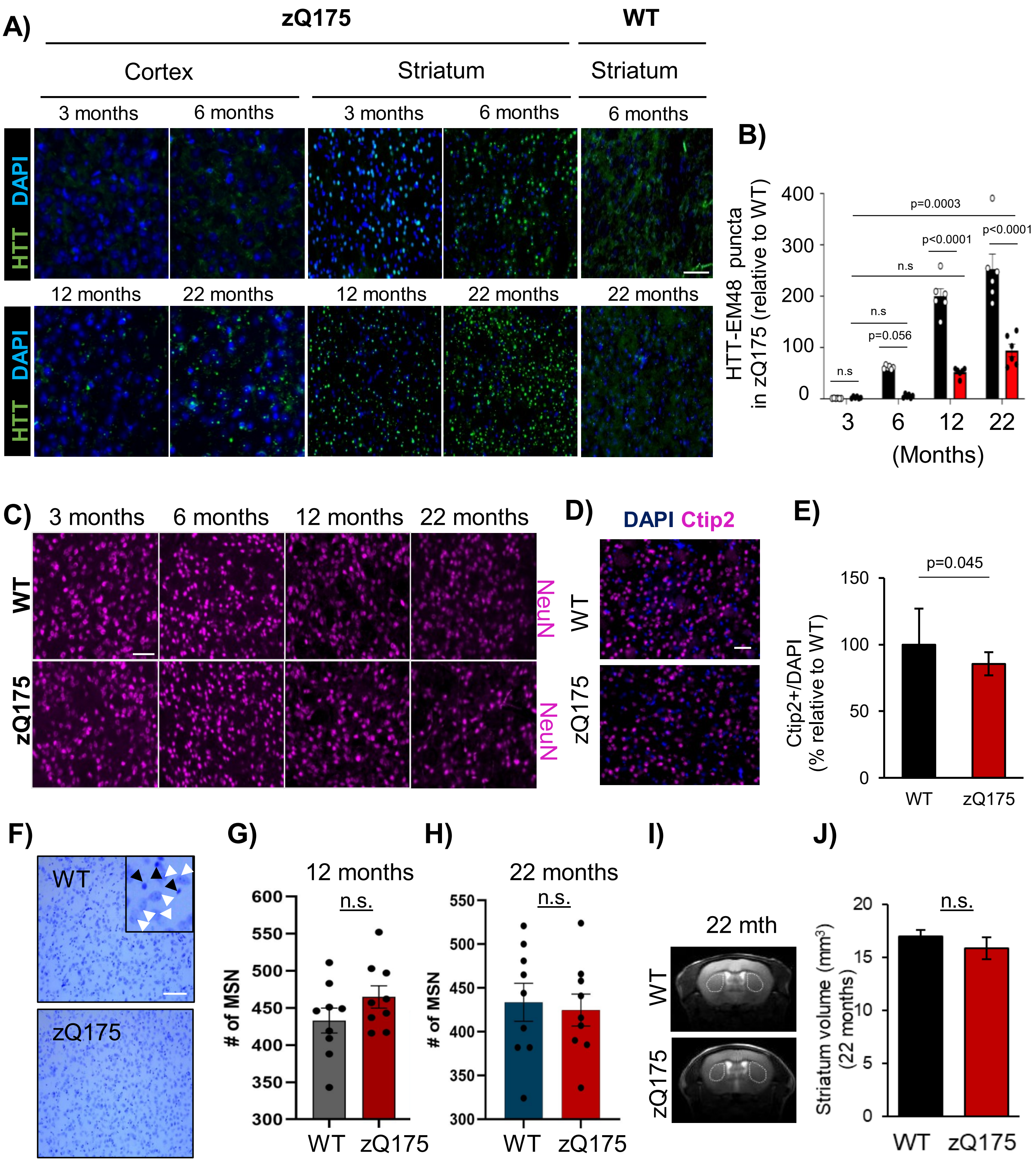

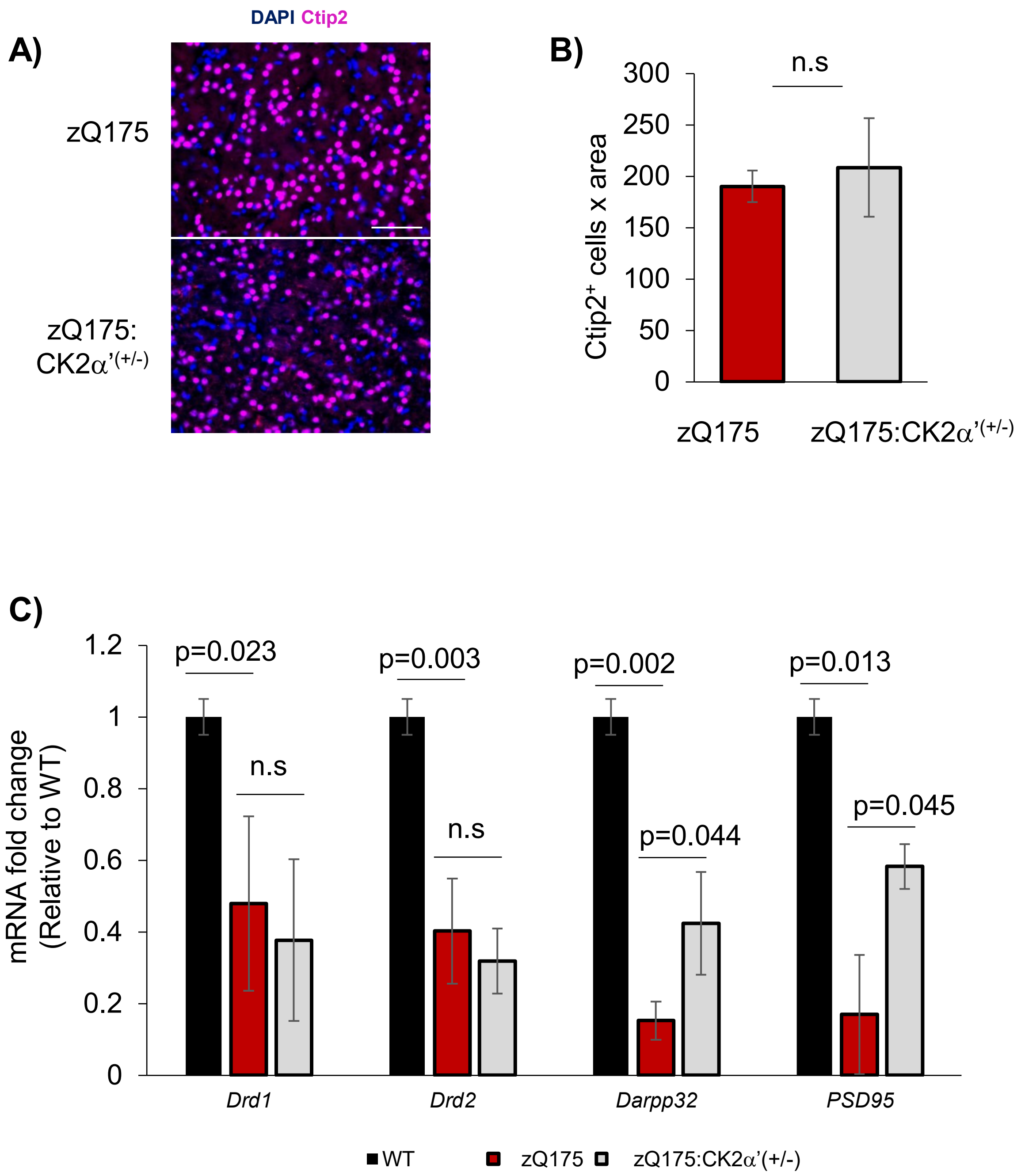

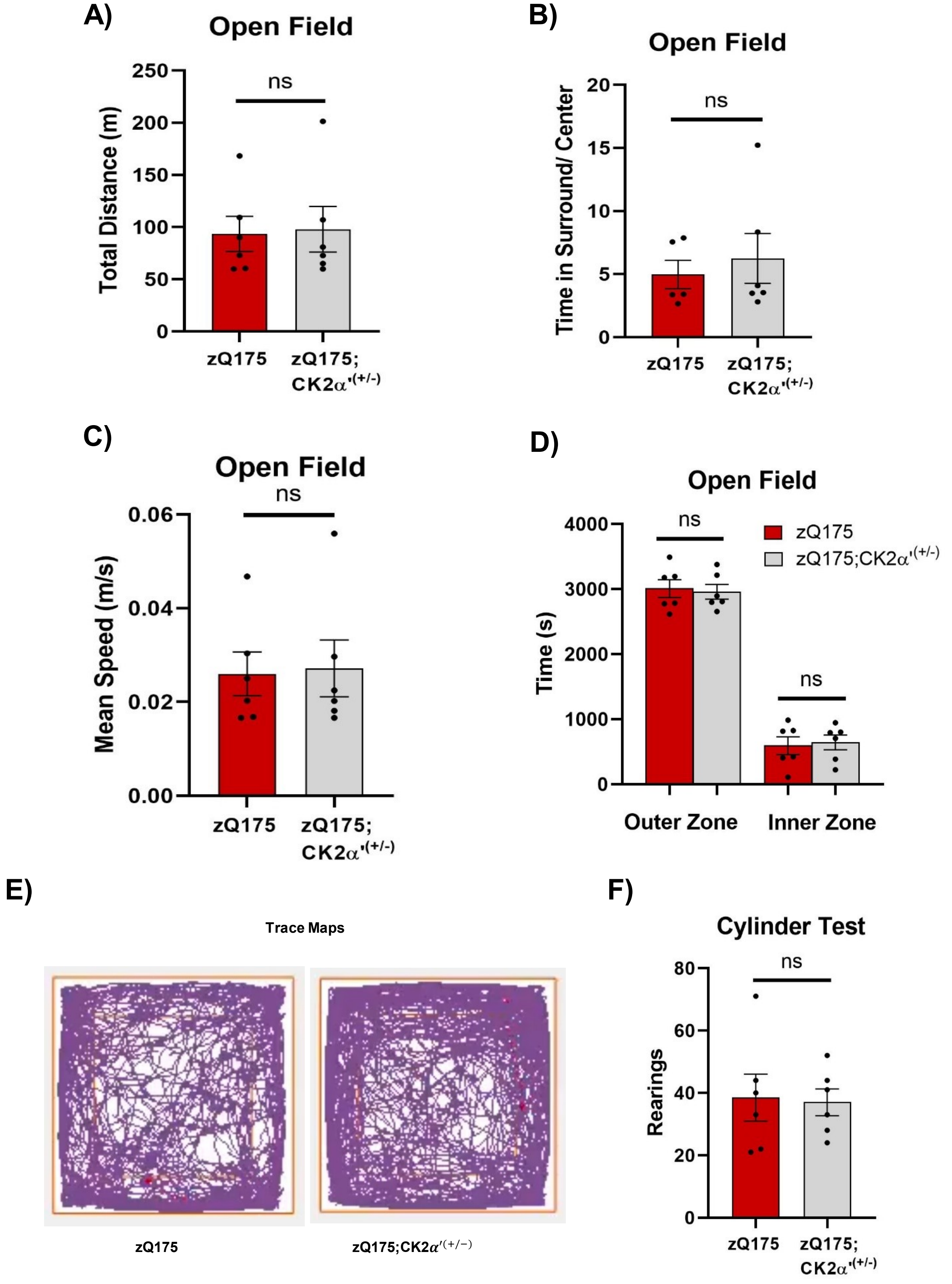

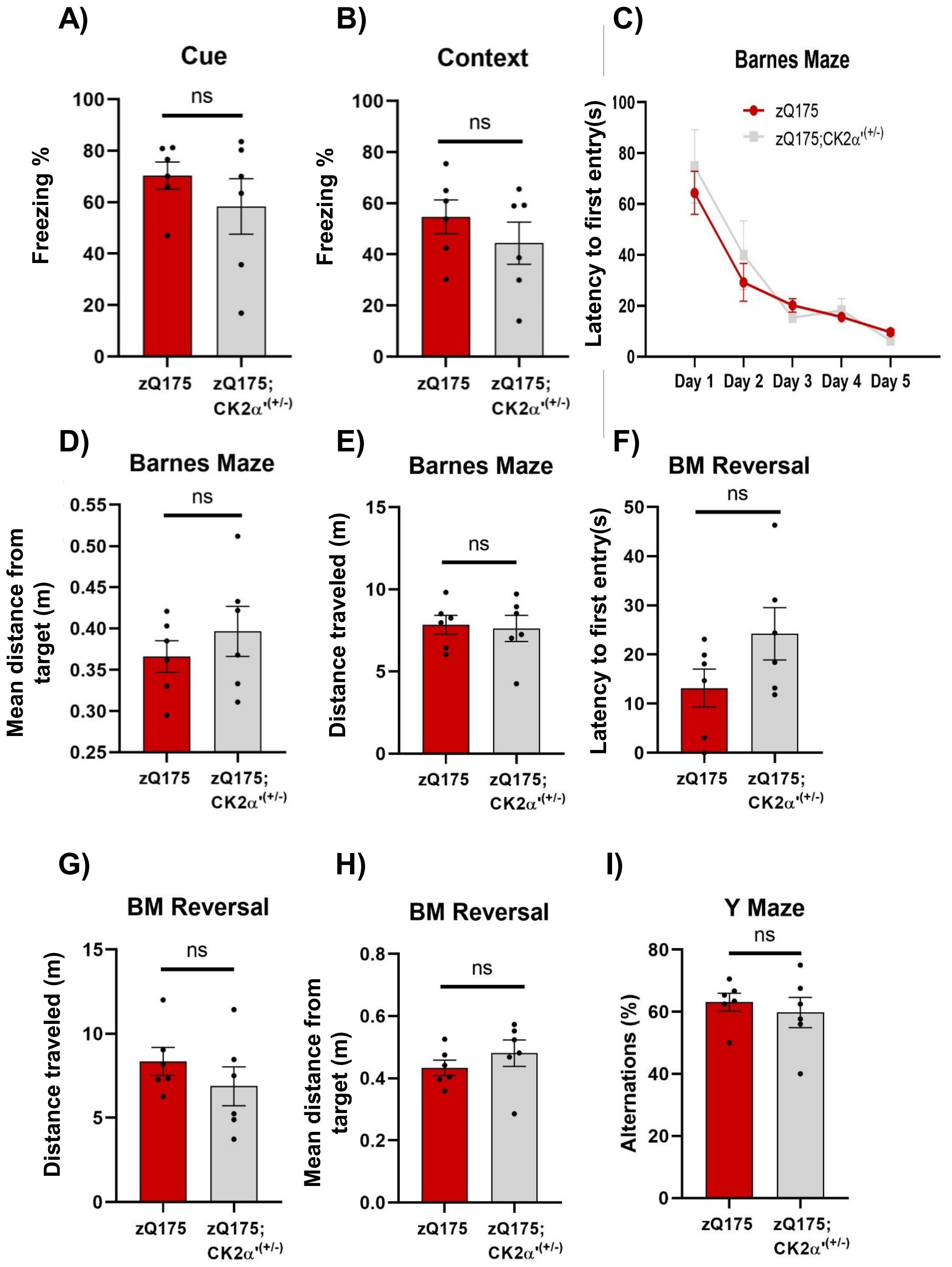

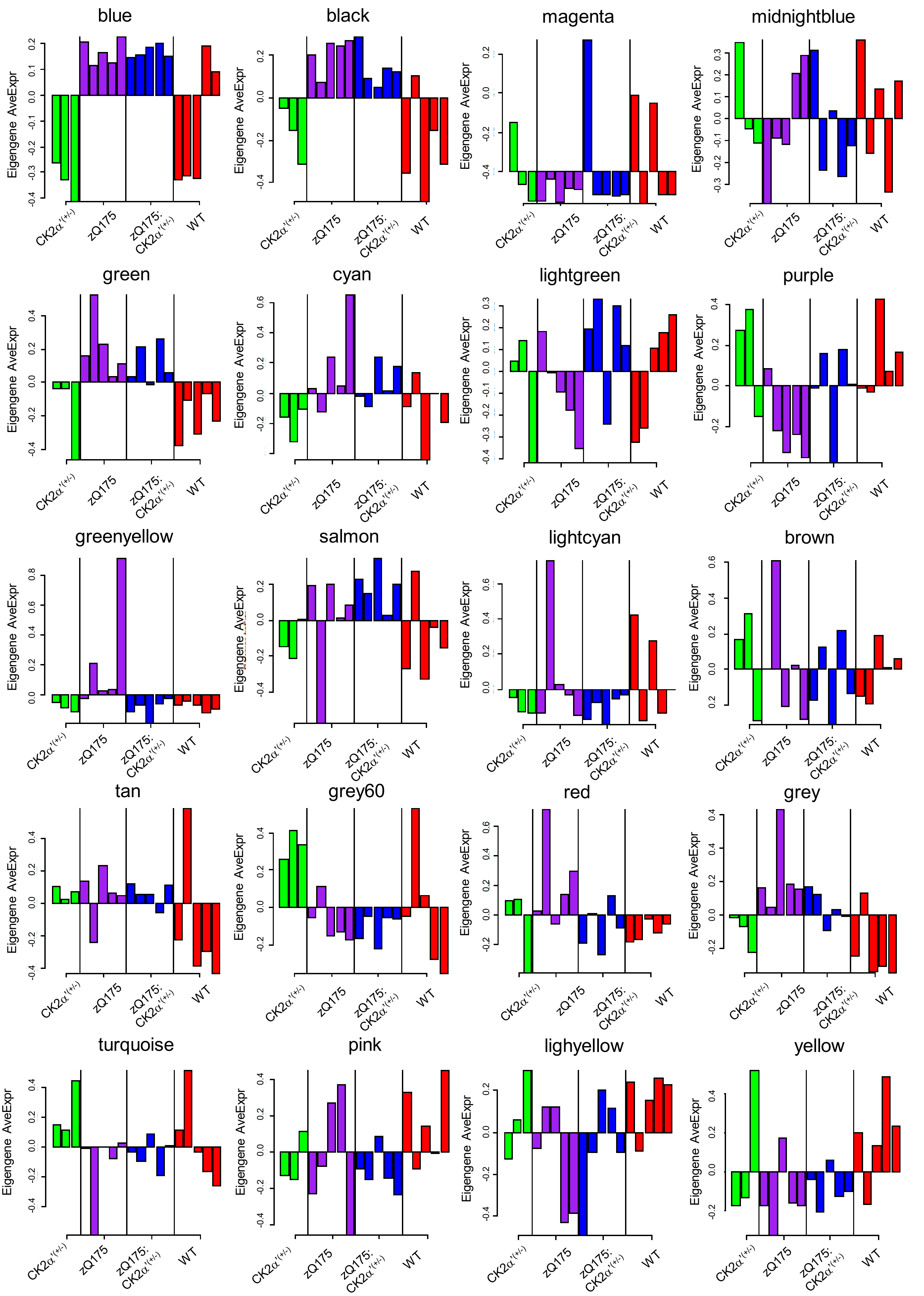

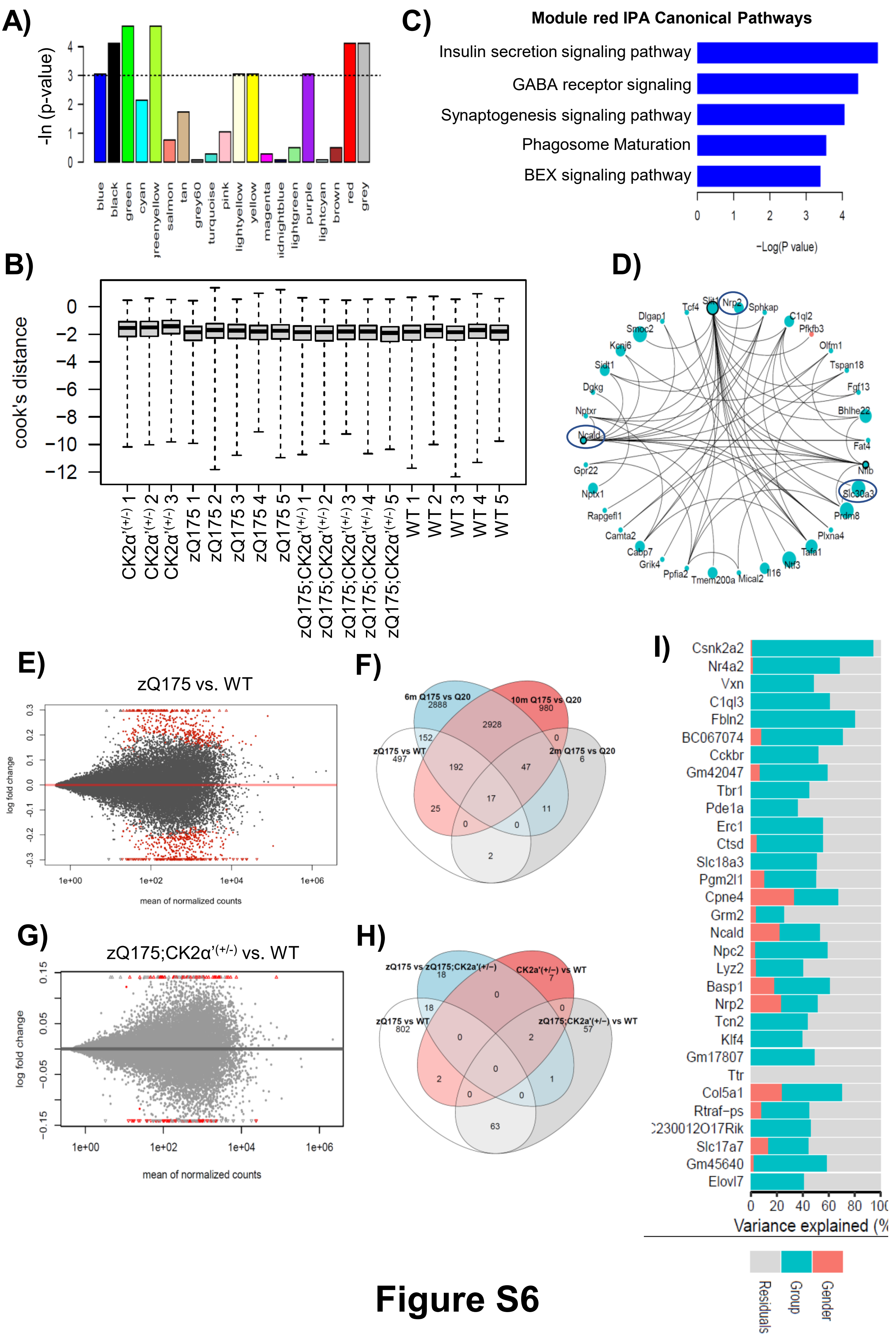

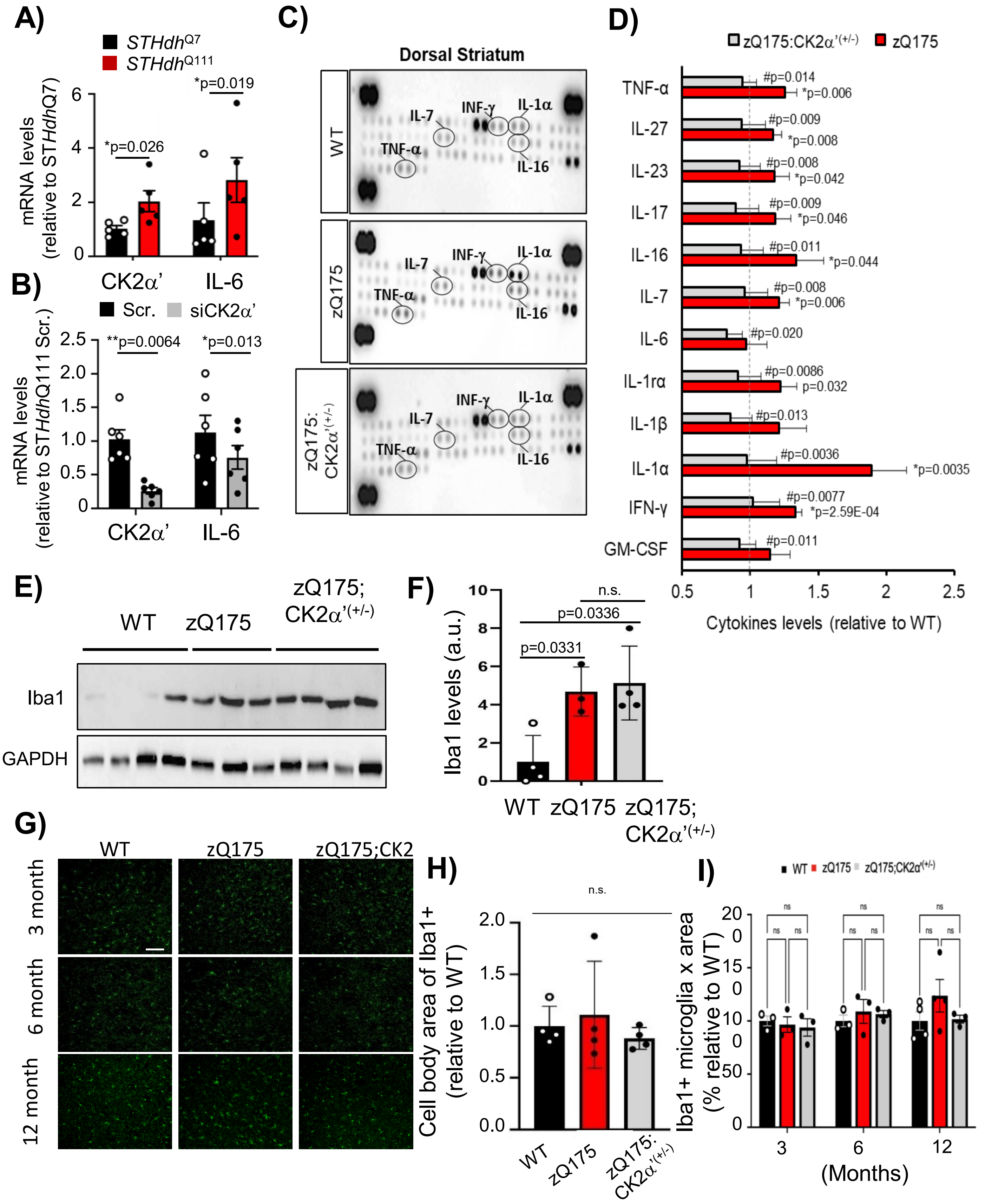

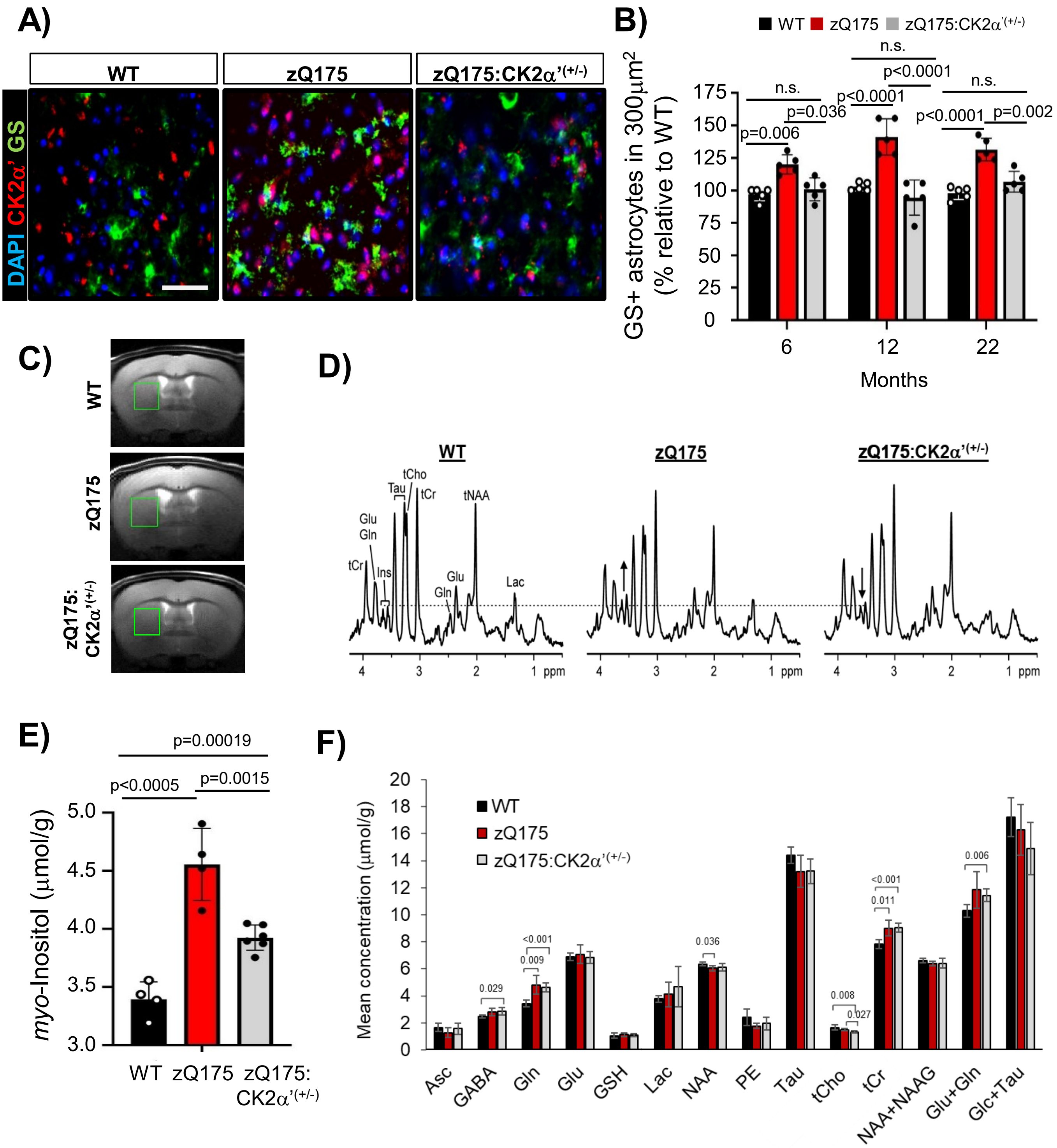

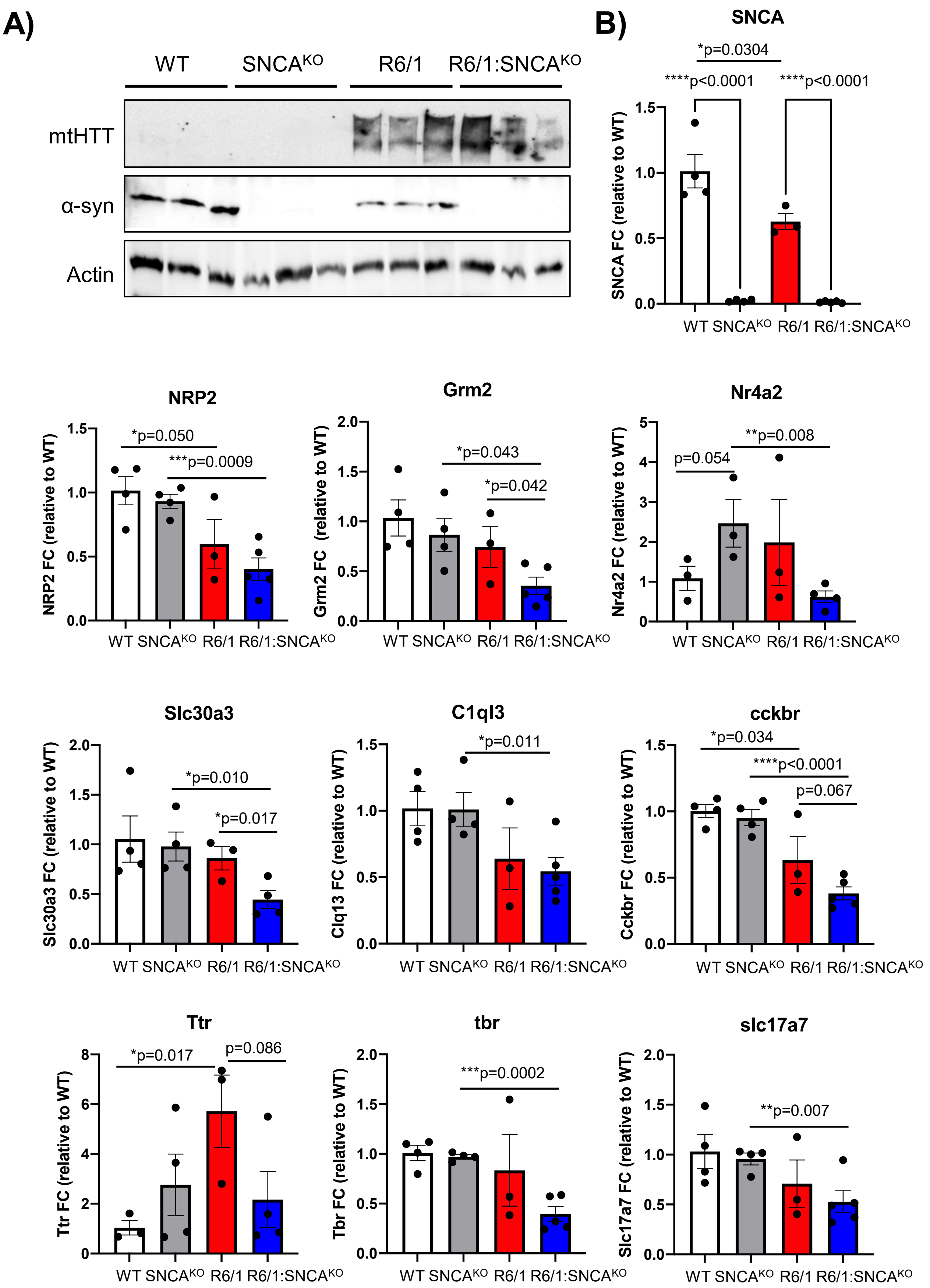

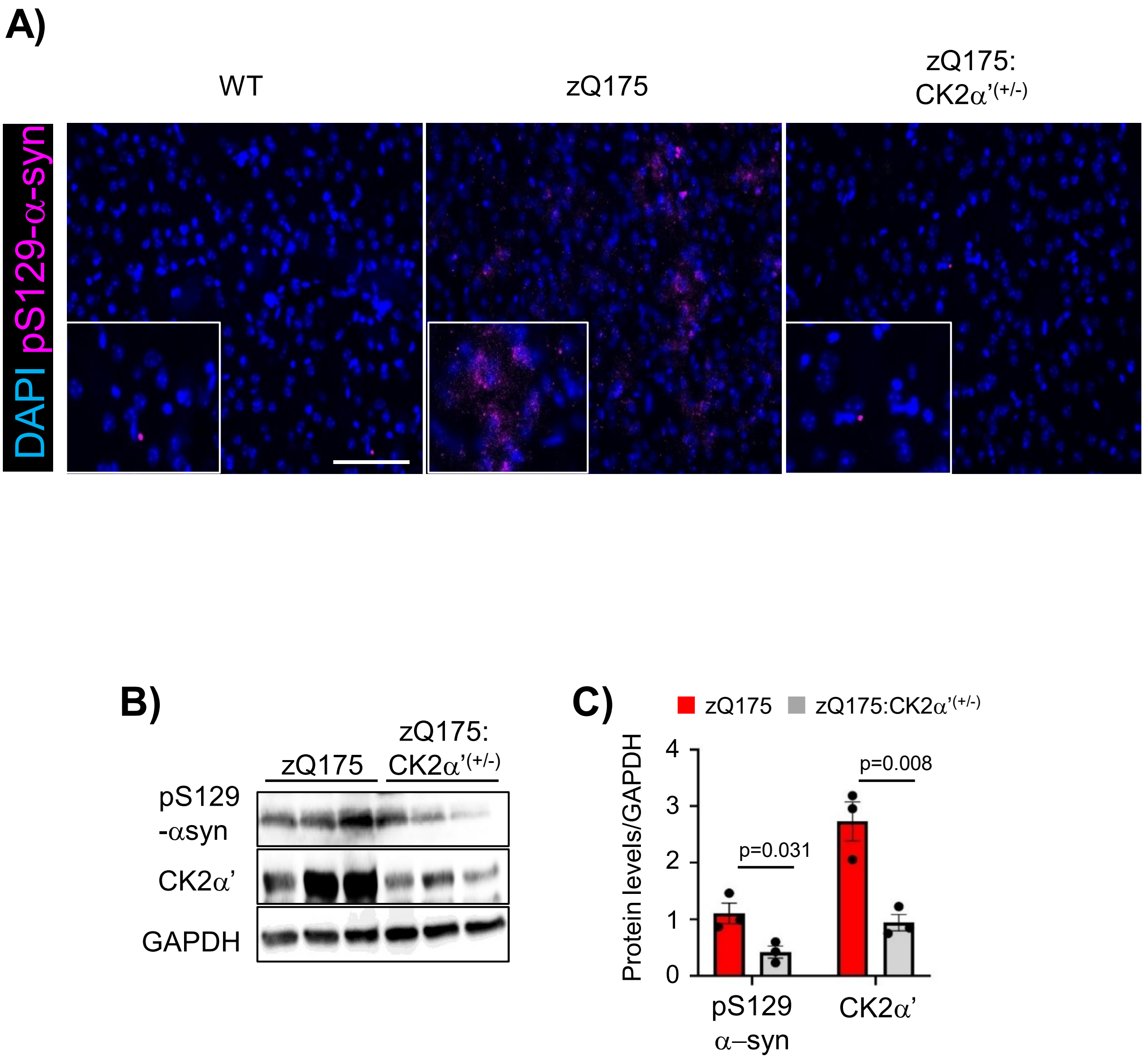
